# Plant-derived soft electrophiles upregulate pro-resolving oxylipins in a paraquat-induced *Drosophila* model of Parkinson’s disease

**DOI:** 10.64898/2026.03.24.714080

**Authors:** Swarnali Chatterjee, Bianca McCarty, Caleb Vandenberg, Madison Bever, Qiaoli Liang, Urmila Maitra, Lukasz Ciesla

## Abstract

Age-accompanied chronic, low-grade systemic inflammation (inflammaging) drives the onset and progression of neurodegenerative disorders like Parkinson’s disease (PD). Currently, no disease-modifying therapies are available for PD. Exposure to environmental toxicants, including paraquat (PQ), rotenone, and neurotoxic metals, increases disease risk. Conversely, sustained consumption of dietary soft electrophiles, such as flavonoids, carotenoids, vitamin E vitamers, and essential fatty acids, has been associated with increased lifespan and delayed age-related neurological decline. Omega-3 and select omega-6 fatty acids also serve as precursors of lipid-derived specialized pro-resolving mediators (SPMs), which exert potent anti-inflammatory and inflammation-resolving activities. Here, we report the development of a robust analytical method to quantify pro-resolving oxylipins in a PQ-induced *Drosophila melanogaster* model of PD, enabling investigation of how dietary phytochemicals modulate anti-inflammatory and pro-resolving lipid metabolism *in vivo*. We hypothesized that plant-derived soft electrophiles promote active resolution of neuroinflammation by enhancing the production of pro-resolving oxylipins derived from essential fatty acids, and that their neuroprotective effects are linked to their soft electrophilic properties. Our results demonstrate that specific lipophilic plant-derived soft electrophiles significantly upregulate pro-resolving oxylipins in *Drosophila* heads following PQ exposure. We identify a subset of flavones and structurally related phytochemicals that selectively enhance SPM biosynthesis and show that this response involves the NF-κB orthologue *relish*. Additionally, feeding modality and sex-specific dimorphisms were found to influence oxylipin production. Collectively, these findings indicate that structurally related dietary soft electrophiles enhance endogenous pro-resolving lipid pathways, promote resolution of toxin-induced neuroinflammation, and have potential preventive and therapeutic relevance for neuroinflammation-associated neurodegenerative diseases.

**Highlights:**

- Quantification of pro-resolving lipids in a *Drosophila* Parkinson’s model.
- Specific structural features of phytochemicals contribute to *in vivo* bioactivity.
- Lipophilic soft electrophiles show therapeutic potential against neuroinflammation.
- Feeding modality and sexual dimorphism also regulate oxylipin production.

**Graphical abstract:** 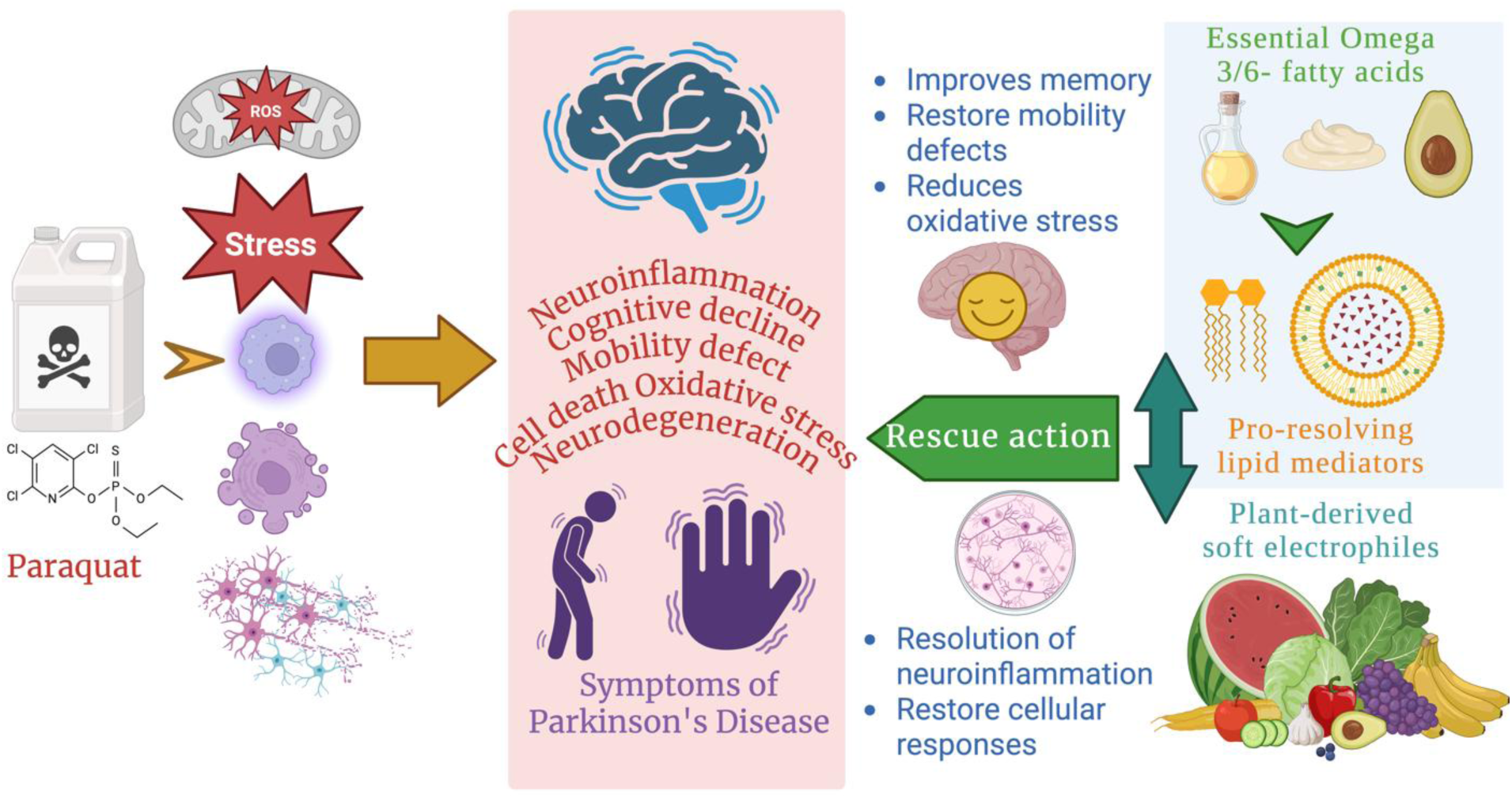

## 1. Introduction

Neurodegenerative diseases constitute a growing global health burden, for which currently available therapies largely provide only symptomatic relief [1]. Among these disorders, Parkinson’s disease (PD) is one of the fastest-growing neurological conditions worldwide [2], particularly in aging populations, with both genetic susceptibility and environmental exposures contributing to disease risk [3]. The herbicide paraquat (PQ) induces oxidative stress, neuroinflammation, motor impairment, and dopaminergic neuron loss and is therefore widely employed as an experimental neurotoxin in diverse PD models [4,5]. Aging itself is accompanied by chronic low-grade systemic inflammation, often referred to as “inflammaging,” which is associated with reduced longevity, cognitive decline, and motor dysfunction [6–8]. Notably, these features overlap with key clinical manifestations of PD [9].

Epidemiological and experimental evidence suggests that regular dietary intake of bioactive compounds, including polyphenols, carotenoids, vitamin E derivatives, essential fatty acids, and particularly flavonoids, may improve cognitive and neurological outcomes in humans [10] and animal models [11]. Polyphenols are abundant plant-derived secondary metabolites commonly consumed as part of dietary patterns such as the Mediterranean diet [12]. Numerous studies associate polyphenol-rich diets with reduced risk of age-related neurodegenerative disorders, including PD, and with attenuation of neuroinflammatory processes [13–15]. Traditionally, the neuroprotective effects of polyphenols have been attributed to their antioxidant, anti-inflammatory, and anti-apoptotic properties [16]. More recent work suggests that these compounds may also modulate gut microbiota, as well as evolutionarily conserved cellular signaling pathways, and thereby influence chronic disease processes [17,18]. Despite substantial supportive evidence, no polyphenolic compounds have yet been approved as prescribed therapeutics [19]. A major limitation is that many plant-derived compounds, particularly flavonoids, are classified as pan-assay interference compounds (PAINS) due to their chemical reactivity and apparent lack of target specificity *in vitro*. As a result, such compounds are frequently deprioritized early in drug discovery pipelines because of false-positive outcomes in high-throughput screening assays [20].

Interestingly, many dietary phytochemicals, including flavonoids, carotenoids, and polyphenols, share structural features consistent with soft electrophilic activity, such as α,β-unsaturated carbonyl groups capable of undergoing Michael addition with nucleophilic cysteine thiols [21,22]. Their biological effects may therefore potentially arise from selective covalent interactions with redox-sensitive protein residues rather than from classical receptor binding. Omega-3 and omega-6 fatty acids (OFAs), as well as their oxidized derivatives known as oxylipins, also possess electrophilic properties [23, 24]. OFAs are essential dietary lipids that support cognition [25], brain function [26], and membrane integrity [27]. Oxylipins derived from OFAs have recently been recognized as pro-resolving lipid mediators that promote tissue repair, restore homeostasis, and actively terminate inflammatory responses [28,29]. Despite their biological importance, oxylipin signaling *in vivo* remains poorly characterized due to their low abundance, rapid turnover, and analytical challenges associated with their detection and quantification [30]. Notably, several established pharmaceuticals, including aspirin, exert part of their anti-inflammatory effects by modulating oxylipin biosynthesis from essential fatty acid precursors and by regulating conserved inflammatory pathways such as Toll/NF-κB and JAK/STAT signaling [31,32].

To investigate the role of oxylipins in neuroinflammation, we employed a paraquat-induced *Drosophila melanogaster* model of PD. *Drosophila* has emerged as a powerful system for studying dietary modulations and neurodegenerative disorders, with approximately 75% of human disease-associated genes having recognizable orthologues in the fly genome [33–35]. PQ-exposed flies recapitulate key features of PD, including oxidative stress, neuroinflammation, and dopaminergic neuron loss [36]. Although lipid metabolism in insects has been investigated, relatively little is known about the presence, regulation, and functional significance of oxylipins in fly neuroinflammatory responses [37,38]. Their low abundance in fly tissues, particularly in the nervous system, has hindered systematic investigation, and robust analytical methods for oxylipin quantification in *Drosophila* are still limited [39]. Consequently, the development of sensitive *in vivo* lipidomic approaches remains essential for advancing our understanding of inflammatory lipid signaling in this model [40].

In the present study, we developed an ultrahigh-performance liquid chromatography–quadrupole time-of-flight high-resolution mass spectrometry (UPLC-QTOF-HRMS) method to identify and quantify two linoleic acid–derived oxylipins, 13-hydroxyoctadecadienoic acid (13-HODE) and 13-oxooctadecadienoic acid (13-oxoODE), in *Drosophila* heads. These metabolites were previously identified as potential anti-inflammatory lipid mediators in flies [31]. Linoleic acid is first converted to 13-HODE and subsequently oxidized to 13-oxoODE (**Fig.1A)**, a soft electrophile implicated in the active resolution of inflammation [31,38]. We established a sensitive and reliable analytical workflow, including a customized extraction protocol optimized for fly head tissue, to quantify these oxylipins following PQ exposure.

**Figure 1:**
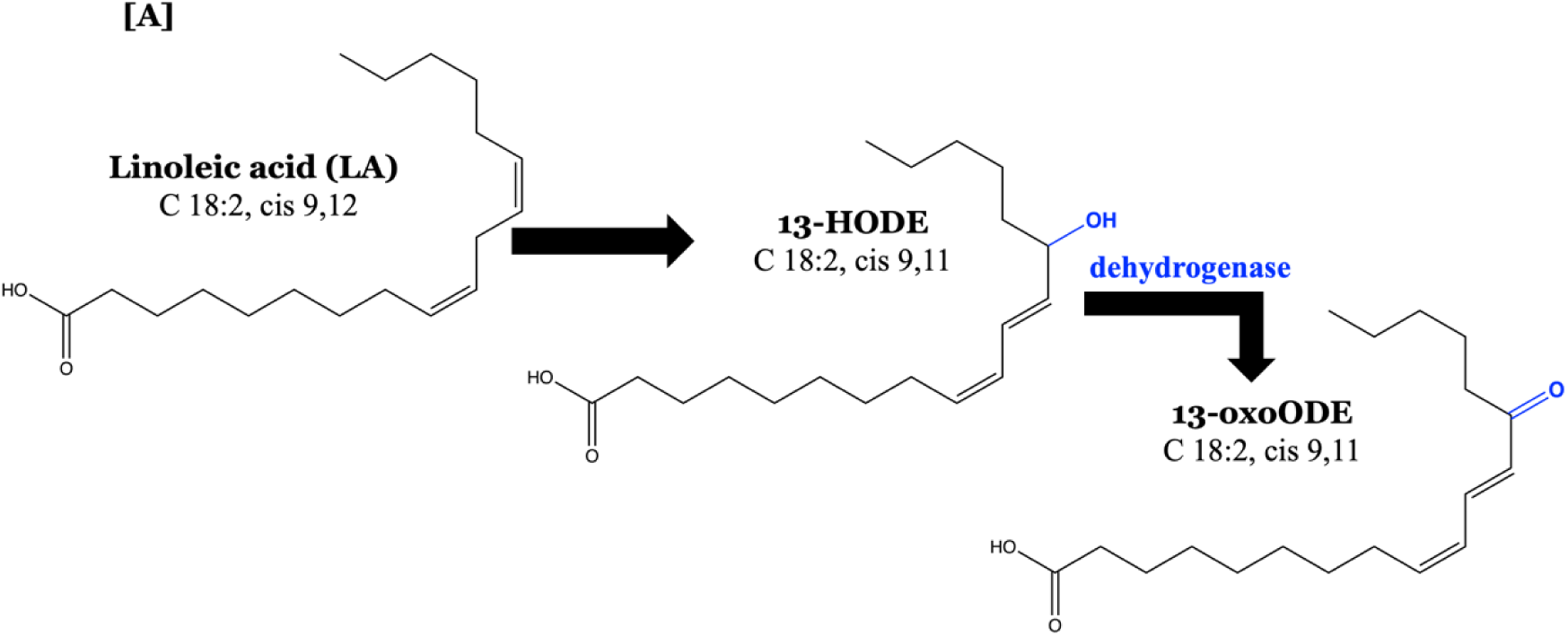
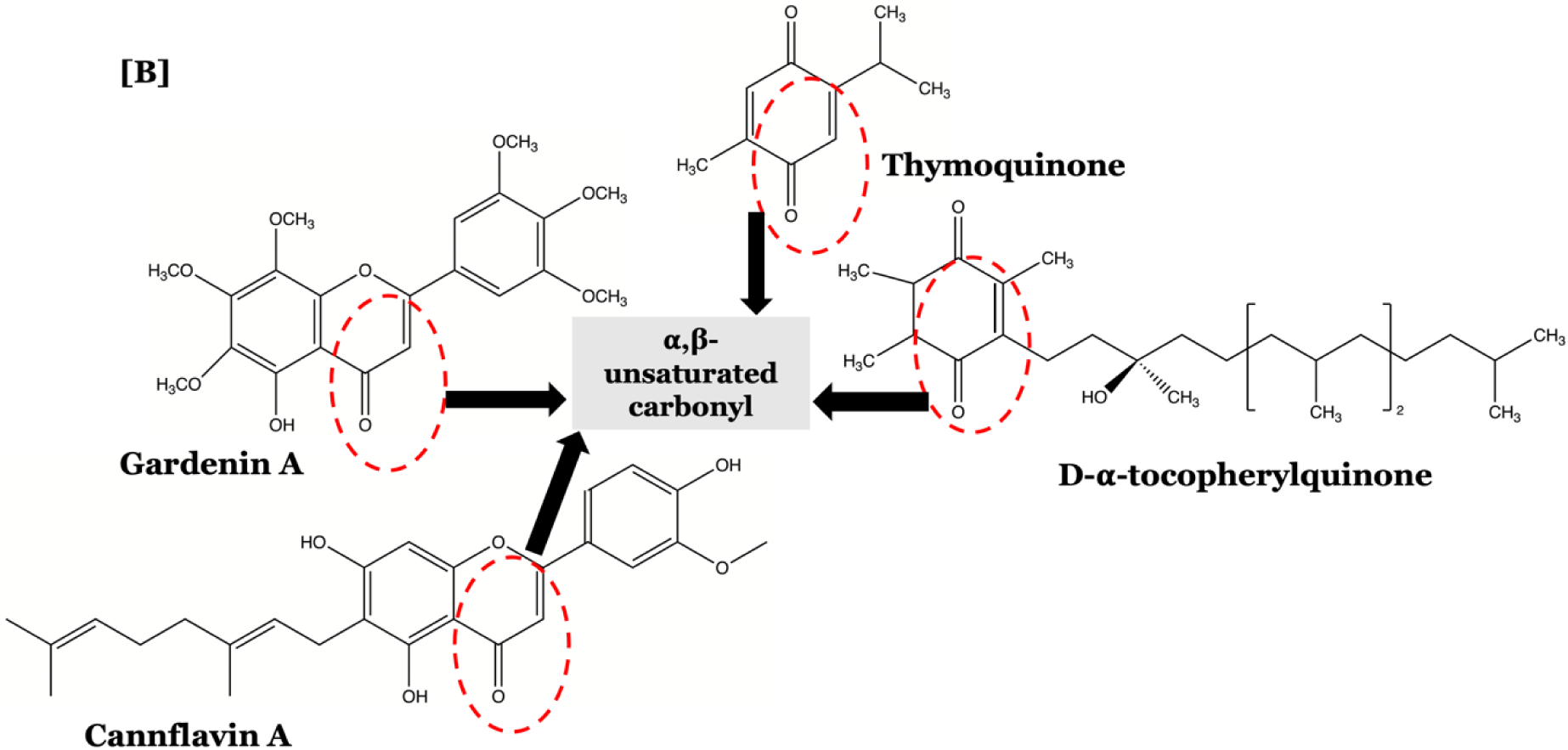
[A] The conversion of precursor Linoleic acid into 13-HODE and 13-oxoODE [31]. [B] The structures of selected plant-derived soft electrophiles indicating the α,β - unsaturated carbonyl group (red circle).

Using this platform, we examined the effects of selected dietary soft electrophiles, including polymethoxyflavonoids, flavones, flavanones, flavonols, quinones, and vitamin E derivatives, on oxylipin production *in vivo*. These compounds were selected based on structural features, like the presence of an α,β -unsaturated carbonyl group associated with soft electrophilic activity; or their ability to generate electrophilic metabolites [41,42]. We observed that pro-resolving oxylipin levels were selectively increased by a specific subclass of plant-derived electrophiles (**Fig.1B**), an effect further enhanced by dietary linoleic acid supplementation [30,31,38]. These findings suggest that both phytochemical structure and lipid availability critically influence oxylipin biosynthesis.

Mechanistic analyses revealed that *relish-null* mutant flies failed to produce anti-inflammatory and pro-resolving oxylipins, implicating conserved NF-κB signaling in oxylipin regulation. This observation aligns with previous evidence demonstrating a central role for NF-κB in coordinating immune and inflammatory responses [6,31,36] and suggests that oxylipin biosynthesis is integrated within canonical inflammatory networks. Furthermore, oxylipin levels were modulated by sex and feeding conditions, with significant differences observed between males and females and across dietary regimens.

In summary, this study introduces a sensitive analytical approach for oxylipin detection in *Drosophila* heads and demonstrates that dietary soft electrophiles can enhance the production of pro-resolving lipid mediators during paraquat-induced neuroinflammation. Our findings support a role for conserved NF-κB signaling in inflammation resolution and reveal complex interactions between diet, sex, and lipid mediator production. Collectively, these results provide a framework for investigating oxylipin biology *in vivo* and underscore the potential of dietary bioactive compounds as modulators of neuroinflammatory processes in PD models.

## 2. Results

### 2.1 Bioactive pro-resolving lipid metabolites are detected in the heads of *wild-type Drosophila* maintained on a linoleic-acid-rich diet post-Paraquat (PQ) exposure

Here, we used a paraquat-induced *Drosophila* model of Parkinson’s disease to detect and quantify bioactive pro-resolving lipid metabolites produced during toxin-induced inflammation, which have previously been observed in *Drosophila* larvae treated with aspirin, a commonly used anti-inflammatory drug [31]. Adult male *Canton S* flies post-eclosion (5 days old) were maintained on diets supplemented with linoleic acid (omega-6 essential fatty acid) in combination with plant-derived soft electrophiles (**Fig.1A**) or control food, then exposed to 5 mM paraquat for 48 hours. Heads were pooled, and lipids extracted for UPLC-QTOF-HRMS analysis. **Figs. 2A and 2B** represent the matrix-matched versus in-solvent (methanol) calibration curves for 13-oxoODE and 13-HODE, respectively. The extracted ion chromatograms (**Fig. 2C**) peak area ratios of 13-HODE [M - H]^−^ *m/z* 295.2273 (RT 8.59 min) and 13-oxoODE [M - H]^−^ *m/z* 293.2117 (RT 8.34 min) versus internal standard ^13^C_18_ linoleic acid [M - H]^−^ *m/z* 297.2928 (10.8 min) were plotted versus their concentration ratios for quantitation. Samples were prepared in triplicate for each concentration, with an average RSD value of 3.61% for 13-oxoODE and 4.15% for 13-HODE. The internal standard normalized standard instrumental matrix effects (ME%) were calculated by the slope ratio of the matrix-matched and solvent calibration curves as 161% for 13-oxoODE and 126% for 13-HODE (**Figs. 2A, 2B**) [43]. The limit of detection (LOD) and limit of quantification (LOQ) were calculated in accordance with the International Conference on Harmonisation (ICH) guidelines. LOD was found to be 0.39 nmol/mL (0.12 ppm) for 13-HODE and 0.36 nmol/mL (0.12 ppm) for 13-oxoODE, and the limit of quantification LOQ was found to be at 1.21 nmol/mL (0.36 ppm) for 13-HODE and 1.1 nmol/mL (0.32 ppm) for 13-oxoODE (refer to **Appendix A. Fig.1** for the specific chromatogram).

**Figure 2:**
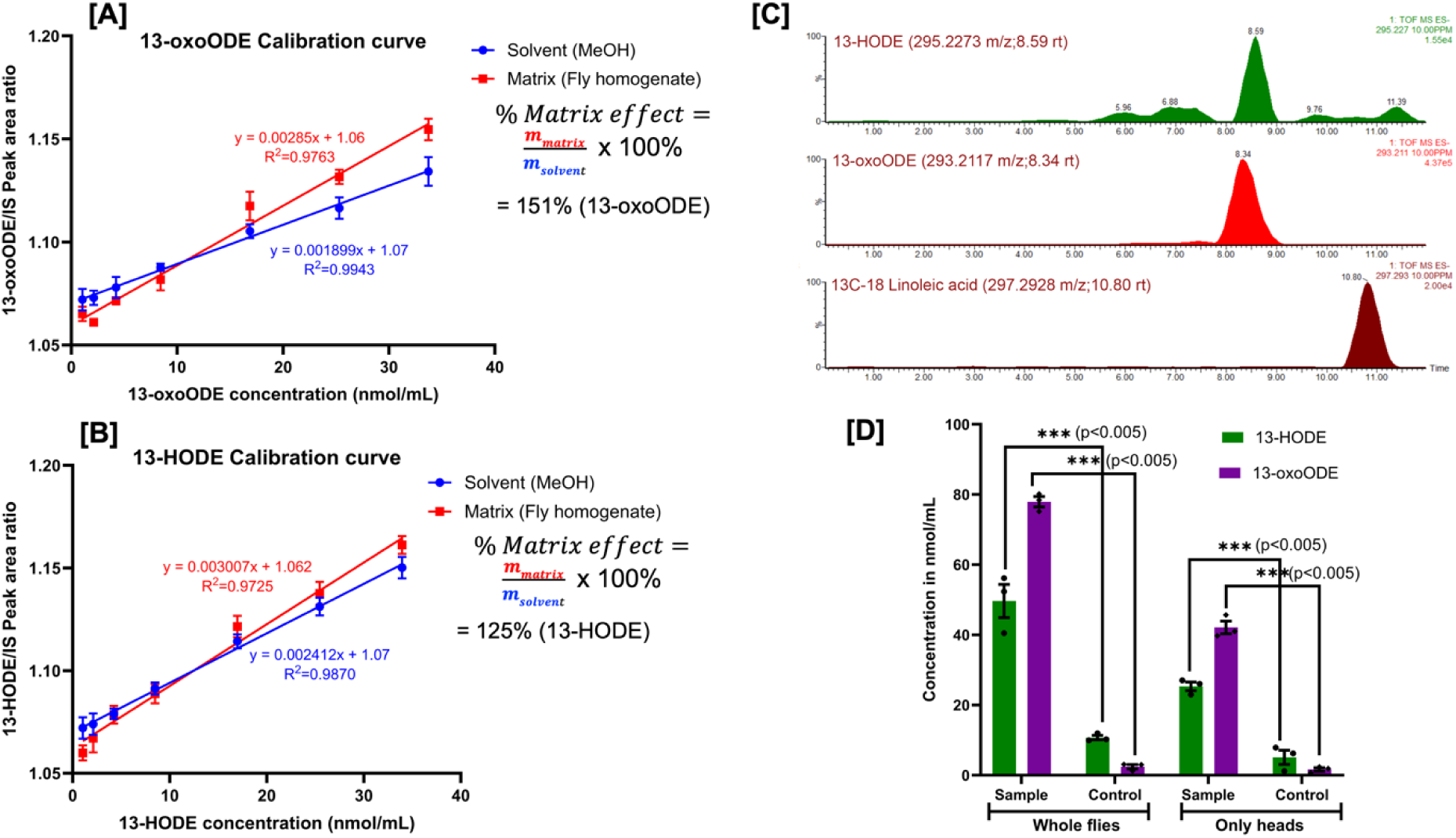
Bioactive pro-resolving lipid metabolites are detected in adult fly heads kept on a linoleic-acid-rich diet post-PQ exposure: <u>Calibration curves for 13-oxoODE [**A**] and 13-HODE [**B**] in solvent (blue) and fly head homogenate (red); HRMS extracted ion chromatograms [**C**] of 13-HODE (*m/z* 295.2273), 13-oxoODE (*m/z* 293.2117), internal standard ^13^C_18_ Linoleic acid (*m/z* 297.2928);</u> and quantification of 13-HODE and 13-oxoODE in *wild type Canton S* adult male fly heads (n=200) v/s whole bodies (n=200) for sample (dietary supplementation of Linoleic acid at 1mM concentration and Gardenin A at 1mM concentration) against control (standard fly diet without any supplementation). Data is representative of three biological and two technical replicates (***p<0.005 represented as mean ± SEM) [**D**].

The concentration of 13-oxoODE was found to be ∼40 nmol/mL in the fly heads vs ∼76 nmol/mL in the whole bodies post-PQ exposure in a homogenate of 50 µL extracted from a group of 200 fly heads. This accounts for about 3.15 ng of 13-oxoODE per fly head and 5.57 ng per whole fly. The data represents at least three biological replicates with at least two technical trials (Mean ± SEM). Internal calibration enabled precise recovery assessment of the internal standard, yielding a recovery rate of 94 ± 1.26% (n=3), demonstrating the robustness and reproducibility of the method. A percent recovery calculation was performed using an internal standard (IS)-based approach. Spiked samples, containing a known concentration of the fly homogenate along with the IS ^13^C_18_ Linoleic acid (^13^C-LA), were analyzed alongside unspiked samples, which contained only the IS and were used to determine the endogenous concentration of the analyte present in the matrix [44]. Additionally, standard solutions with known concentrations of both 13-HODE, 13-oxoODE and IS were analyzed to establish a calibration curve for quantification. This calculation allowed for the evaluation of extraction efficiency and method performance. This validated approach was employed for the relative quantification of 13-HODE and 13-oxoODE in *wild-type Canton S* adult male *Drosophila* tissues, comparing lipid concentrations in isolated heads versus whole bodies following PQ exposure.

Our results demonstrate that in response to dietary supplementation with plant-derived soft electrophiles, followed by exposure to PQ for 48 hours, *Drosophila* produces pro-resolving lipid mediators, namely 13-HODE and 13-oxoODE. To the best of our knowledge, this is the first evidence of the quantification of bioactive lipid pro-resolving molecules in *Drosophila* heads. After PQ treatment, we found that dietary intervention with linoleic acid supplementation, combined with PQ-induced oxidative stress, significantly enhanced the synthesis of 13-oxoODE, the final oxidized metabolite derived from linoleic acid.

### 2.2 Selected plant-derived soft electrophiles increase survival, improve mobility, and upregulate oxylipin production in *wild-type Drosophila* heads post-PQ exposure

In this study, adult male *Drosophila (wild-type Canton-S)*, aged five days post-eclosion, were treated with linoleic acid and plant-derived soft electrophiles. Specific pretreatments included each of Gardenin A, Thymoquinone, D-alpha-tocopherylquinone (DATQ), Apigenin, Nobiletin, Cannflavin A, and Puresirtmax, in combination with linoleic acid, followed by PQ exposure. These compounds were evaluated for their protective effects against toxicity induced by 5 mM Paraquat (PQ) exposure. Survival assays were conducted by counting live flies every 24 hours until death, demonstrating that the plant-derived soft electrophiles protected the flies against PQ-induced toxicity by significantly increasing their lifespan. In addition, negative geotaxis assays were used to assess locomotor function under PQ-induced toxicity, measuring the climbing ability of flies at 24- and 48-hours post-PQ treatment. After rigorous preliminary screening, it was observed that equal concentrations of both the phytochemical and linoleic acid worked as the best combination to produce oxylipins in the flies that were in a detectable and quantifiable range. Thus, the standard fly food was supplemented with each phytochemical and linoleic acid at a final concentration of 1mM. Efficient food preparation was performed to ensure that the test compounds were thoroughly mixed into the standard fly food and that the flies consistently consumed the modified diet. To ensure that each fly received the same volume of food during the 5-day feeding period, the control diet contained an equal volume of 2.5% sucrose in place of the phytochemical mixture. This sucrose solution was also used to dilute the stock solutions of the soft electrophiles. The feeding method and phytochemical dosages were validated in preliminary survival assays, which identified the minimum millimolar concentrations that were non-lethal and produced the strongest protective effects. The food uptake was confirmed by detecting the peak of linoleic acid in the fly head homogenate using UPLC-HRMS since linoleic acid has been used to enrich the standard fly food in both the sample and control groups (**Appendix A. Fig. 2**). Previous studies have demonstrated that Gardenin A, Apigenin, and Nobiletin significantly increase lifespan and improve the climbing index in Canton-S adult male flies following PQ exposure [36,41]. Consistent with these findings, **Figs. 3A and 3B** show that Thymoquinone, D-alpha-tocopherylquinone, Cannflavin A, and Puresirtmax exhibit similar activity. The observed effects suggest that these compounds, which share structural similarities with Gardenin A, Apigenin, and Nobiletin, confer protective benefits by improving lifespan and locomotor performance in flies after PQ exposure.

**Figure 3:**
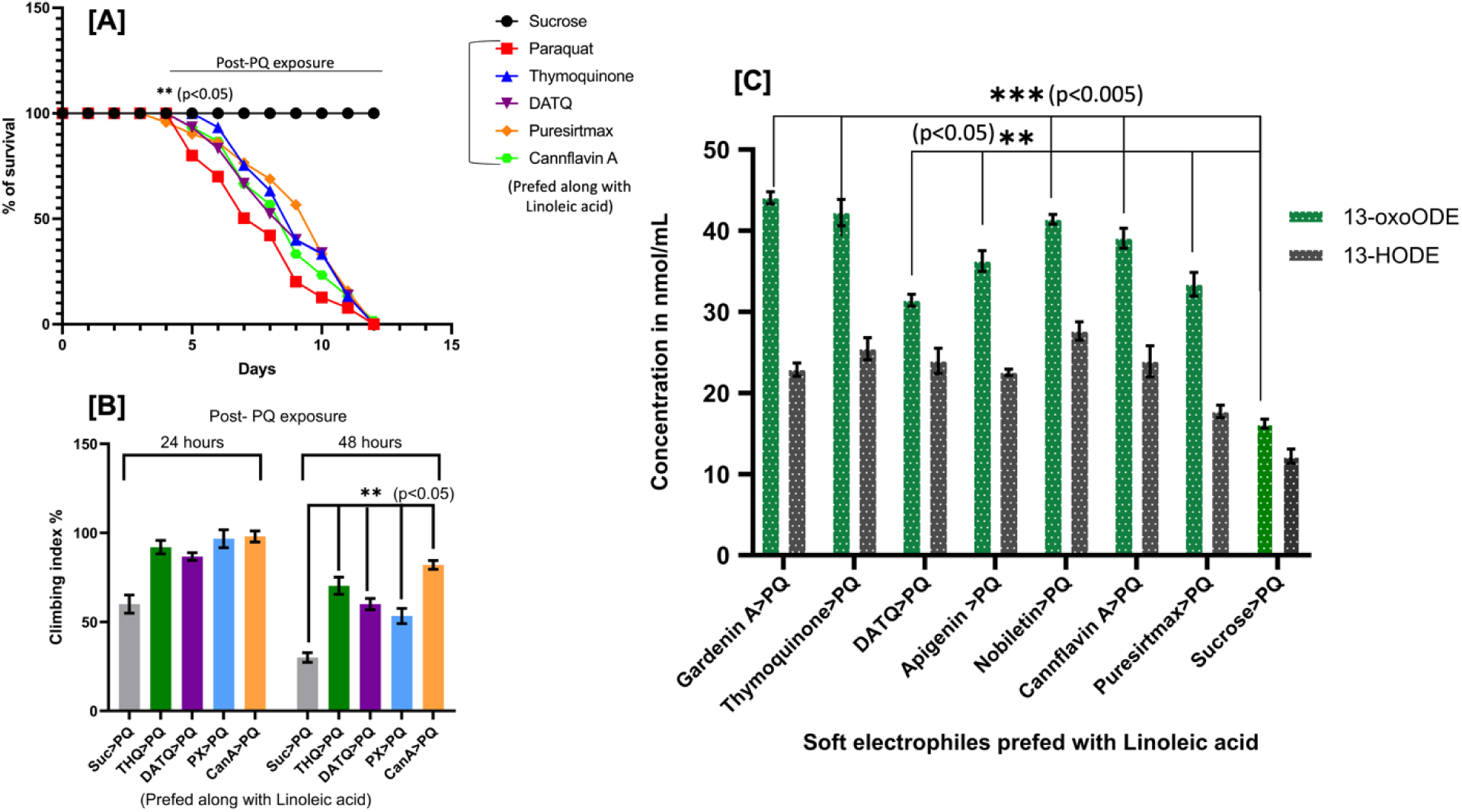
Selected plant-derived soft electrophiles upregulate oxylipin production in wild-type fly heads post-PQ exposure: **[A]** <u>Survival curve representing the protective effects of dietary supplementation of linoleic acid and specific plant-derived soft electrophiles against PQ-induced toxicity.</u> 10 separate biological replicates with 10 flies per feeding condition are shown in the data. The data displayed is mean ± SEM. **p<0.05 according to the Kaplan-Meier survival test. **[B]** <u>Negative geotaxis assay is represented as a measure of climbing index 24hrs and 48hrs post-PQ exposure.</u> 10 separate biological replicates with 10 flies per feeding condition are shown in the data. The data displayed is mean ± SEM. **p<0.05 according to one-way ANOVA between the specific dietary supplementation with the particular soft electrophiles (Fig.1A). **[C]** <u>The graph represents a ***relative upregulation*** of 13-oxoODE and 13-HODE (</u>Fig 1B<u>) in *wild type Canton S* adult male fly heads (n=200)</u>. The concentration of 13-oxoODE was 44.136 nmol/mL in the Gardenin A fed fly heads, 41.072 nmol/mL in the Thymoquinone fed fly heads as compared to standard fly food (control group) post-PQ exposure. The data represent at least three biological replicates with at least two identical trials (Mean±SEM, *\*\*\**p<0.005*, \*\**p<0.05 for one-way ANOVA).

Furthermore, targeted lipidomic analysis revealed a relative upregulation of the linoleic acid-derived pro-resolving metabolites 13-oxoODE and 13-HODE in the heads of pretreated flies. To accurately measure these oxylipins, which are produced in low abundance in *Drosophila*, we developed and employed a tailored analytical method optimized for fly samples. Using this approach, both 13-HODE and 13-oxoODE were successfully detected and quantified in fly head homogenates with the aid of the isotopically labeled internal standard ^13^C_18_-linoleic acid. The total analytical run time for each sample was maintained at 14 minutes.

Quantification was performed using internal-standard peak area ratios, calculated as the ratio of the area under the sample peak to that of the corresponding internal standard peak. This relative area ratio was then converted to concentration using the calibration curve, enabling accurate quantification of the target lipid metabolites. Using this method, concentrations of 13-oxoODE reached 44.136 nmol/mL (3.235 ng/head) in Gardenin A–fed flies and 41.072 nmol/mL (3.011 ng/head) in Thymoquinone-fed flies. These levels represent an approximately 3–4-fold increase compared with sucrose-only controls (10.19 nmol/mL; 1.18 ng/head) following PQ exposure, as shown in Fig. 3C (n ≥ 3 biological replicates; ***p < 0.0005, **p < 0.005; one-way ANOVA).

Collectively, these results indicate that soft electrophilic phytochemicals, including Thymoquinone, Gardenin A, D-alpha-tocopherylquinone, Apigenin, Nobiletin, Cannflavin A, and Puresirtmax promote the upregulation of oxylipin production in adult male *Drosophila* heads.

### 2.3 Loss of *Relish*, the human NF-kB orthologue in *Drosophila*, disrupts the production of pro-resolving oxylipins post-PQ exposure

Relish null mutant adult male flies (5 days post-emergence) were maintained on a diet supplemented with plant-derived soft electrophiles and linoleic acid. Survival was monitored by counting the number of live flies every 24 hours until all flies had died, and survival percentages were plotted over time to generate the survival curves shown in **Fig. 4A**. The dataset represents 10 independent biological replicates, each consisting of 10 flies per feeding condition. Kaplan-Meier survival analysis revealed no statistically significant differences among the treatment groups, indicating that the tested plant-derived soft electrophiles did not affect the lifespan of relish null mutant adult male flies following PQ exposure.

**Figure 4:**
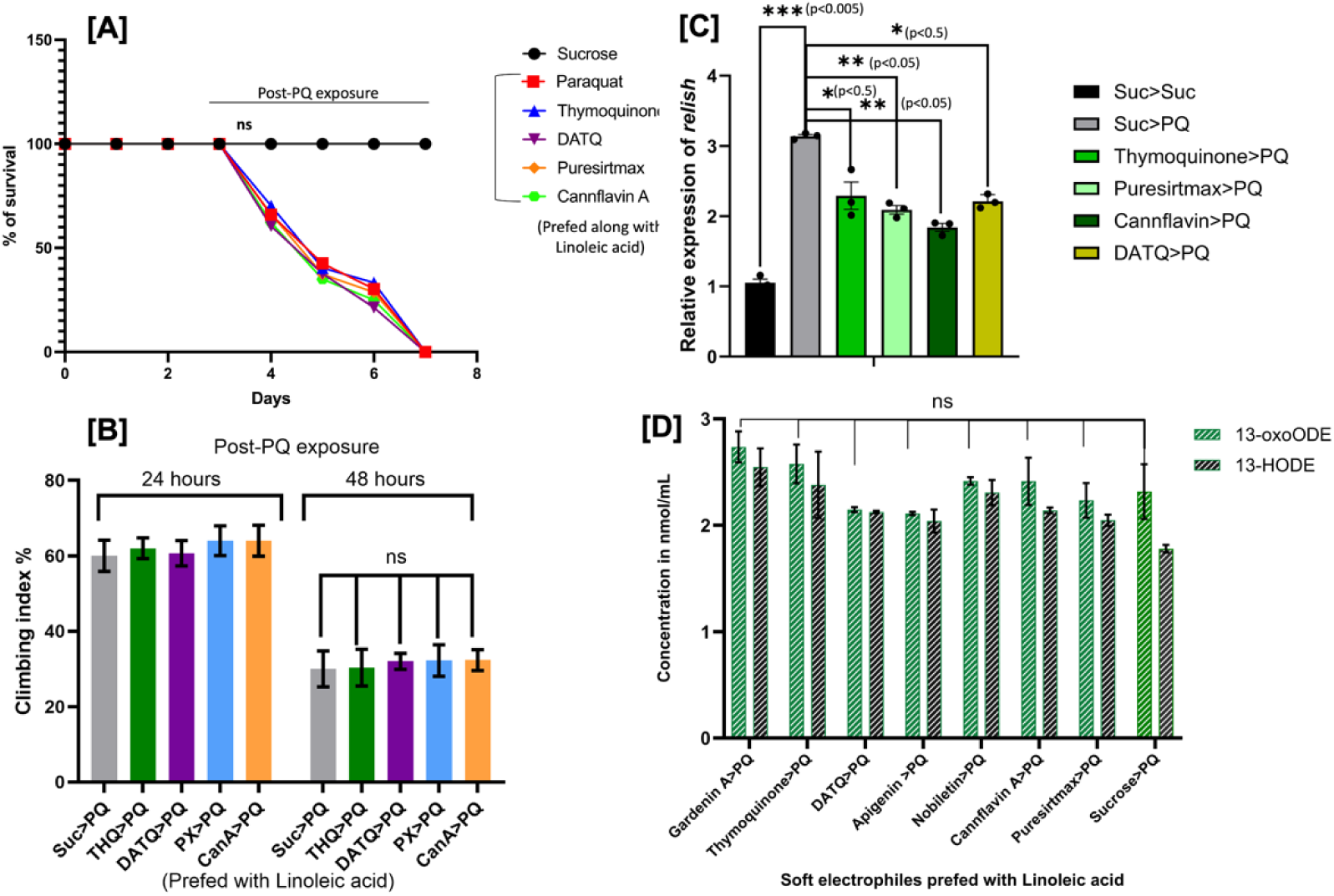
Loss of Relish, the human NF-kB orthologue in Drosophila, disrupts the production of pro-resolving oxylipins post-PQ exposure: **[A]** <u>Survival curve representing the effects of dietary supplementation of linoleic acid and specific plant-derived soft electrophiles against PQ-induced toxicity in *relish null mutant* adult male flies.</u> 10 separate biological replicates with 10 flies per feeding condition are shown in the data. *ns* represents that the data was not significant between the control group (Suc>PQ) and the different feeding conditions that were considered according to the Kaplan-Meier survival test. **[B]** <u>Negative geotaxis assay is represented as a measure of climbing index 24hrs and 48hrs post-PQ exposure.</u> 10 separate biological replicates with 10 flies per feeding condition are shown in the data. The data displayed is mean ± SEM. *ns* represents that the data was not significant between the control group and the different feeding conditions that were considered according to one-way ANOVA. **[C]** <u>The graph represents relative effects of Thymoquinone, Puresirtmax, Cannflavin A and DATQ on the transcript levels of *relish* in *wild-type Canton S male adult flies*, post-PQ-exposure, using qRT-PCR</u>. *Relish* transcript levels were analyzed and plotted after normalization to *rp49* levels as the internal control. Each data point represents mean ± SEM. The mRNA fold changes are normalized to sucrose-fed flies (Suc>Suc; assigned a value of 1). *p<0.5, **p<0.05, ***p<0.005 were considered significant according to the Mann-Whitney U test between Suc>PQ and the different feeding conditions considered. **[D]** <u>The graph represents relative quantities of 13-oxoODE and 13-HODE in *relish null mutant* adult male fly heads (n=200).</u> The concentration of 13-oxoODE was 2.813 nmol/mL in the Gardenin A fed fly heads, 2.622 nmol/mL in the Thymoquinone fed fly heads which is almost equivalent to the average oxylipin production in the 2.5% sucrose-fed fly heads (control) post-PQ exposure.

To assess locomotor performance, negative geotaxis assays were conducted to determine whether pretreatment influenced the climbing ability of flies exposed to PQ (**Fig. 4B**). As with the survival assay, 10 independent biological replicates with 10 flies per feeding condition were analyzed. One-way ANOVA revealed no statistically significant differences among the dietary supplementation groups, indicating that the tested plant-derived soft electrophiles did not improve locomotor function in relish-null mutant adult male flies following PQ exposure.

Given the lack of phenotypic effects on lifespan and mobility, we next examined whether these phytochemicals influence the expression of relish, the Drosophila orthologue of human NF-κB. Quantitative RT-PCR analysis was performed on RNA extracted from fly heads after exposure to the designated feeding conditions. Relish transcript levels were normalized using rp49 as the internal control, and flies maintained on sucrose alone (Suc > Suc) were used as the baseline reference (assigned a value of 1). Notably, all tested phytochemicals significantly suppressed relish transcript expression (Fig. 4C).

Previous studies demonstrated that Gardenin A, Apigenin, and Nobiletin significantly prolonged lifespan, improved climbing ability, and reduced relish transcript levels in PQ-exposed Canton S adult male flies [36,41]. Consistent with these findings, **Figs. 3A and 3B** show that Thymoquinone, D-alpha-tocopherylquinone, Cannflavin A, and Puresirtmax, compounds that share structural similarities with Gardenin A, Apigenin, and Nobiletin exert protective effects on lifespan and climbing performance following PQ exposure. In addition, **Fig. 4C** shows that these compounds significantly reduced relish transcript levels by approximately 30–50%, consistent with our previous findings on structurally related polymethoxy flavonoids [41].

To further investigate the role of relish in inflammatory resolution, targeted lipidomic analysis was performed to determine whether the absence of relish influences oxylipin production following PQ-induced inflammation. Interestingly, no significant upregulation of 13-oxoODE was observed in relish-null mutant flies compared with the control group (**Fig. 4D**). The average levels of both 13-HODE and 13-oxoODE in relish null mutants were comparable to those of controls. Specifically, the quantities of 13-oxoODE were approximately 2.81 nmol/mL in Gardenin A-supplemented flies and 2.57 nmol/mL in Thymoquinone-supplemented flies, representing nearly a 15-fold decrease compared with wild-type Canton S adult male flies. Quantification was performed using internal standard peak area ratios, calculated as the ratio of the area under the sample peak to that of the corresponding internal standard peak.

A similar trend was observed in relish-null mutant adult female flies, with no significant differences relative to males (data not shown). Collectively, these findings strongly suggest that relish plays a critical role in the regulation of pro-resolving oxylipins. Given that relish is the *Drosophila* orthologue of human NF-κB, these results further support the evolutionary conservation of inflammatory signaling and resolution mechanisms between Drosophila and mammals.

### 2.4. Non-neuroprotective flavonoids do not upregulate the production of pro-resolving oxylipins post-PQ exposure

A group of structurally related flavonoids, including Quercetin (**Fig. 5B**), Gardenin B, Myricetin, Fisetin, Hesperetin, and Naringin, previously reported to lack neuroprotective effects were administered to wild-type Canton S adult male flies together with linoleic acid through standard fly food. Although lipophilic neuroprotective flavones and anti-inflammatory lipid-derived oxidation products of essential fatty acids share certain structural characteristics, flavonoids within this group differ in key structural features that define major flavonoid subclasses and influence their biological activity. In particular, variations in hydroxylation or methoxylation patterns and the degree of unsaturation within the C-ring are known to modulate flavonoid bioactivity [36,41,65]. Structural analysis of these molecules revealed notable differences in functional group distribution. For example, Quercetin, Myricetin, and Fisetin are characterized by the presence of a hydroxyl group at the C-3 position of the flavonoid C-ring along with an α,β-unsaturated carbonyl moiety. In contrast, Hesperetin and Naringin lack these structural characteristics (structures shown in **Appendix A, Fig. 4**).

**Figure 5:**
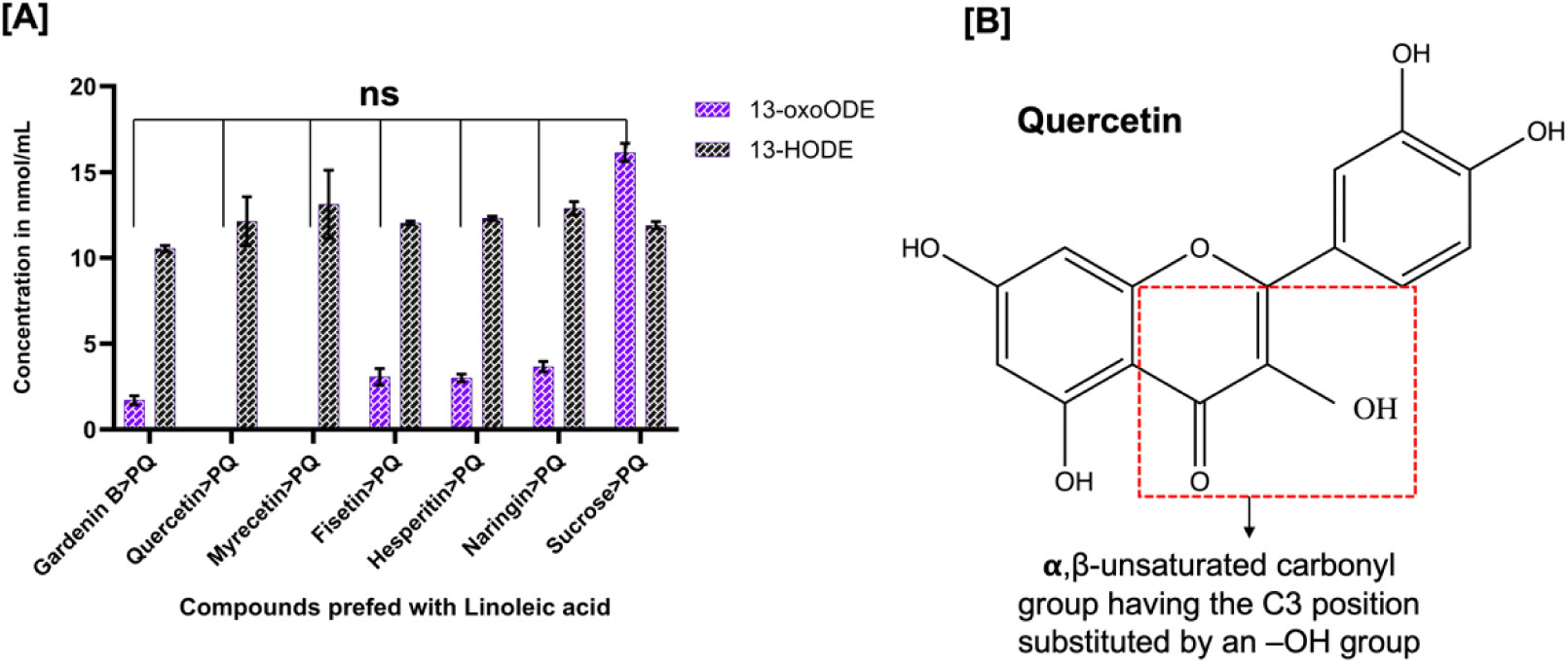
Non-neuroprotective flavonoids do not upregulate the production of pro-resolving oxylipins post-PQ exposure: **[A]** <u>The graph represents relative quantities of 13-oxoODE and 13-HODE in *wild type Canton S* male fly heads (n=200).</u> The concentration of 13-oxoODE was 2.361nmol/mL (0.173 ng/head) in the Hesperitin-fed fly heads, 1.893 nmol/mL (0.139 ng/head) in the Gardenin B-fed fly heads which is almost equivalent to the average oxylipin production in the 2.5% Sucrose fed fly heads (control) post-PQ exposure (n=200). **[B]** <u>The structure of quercetin represents the substitution of C3-position with a hydroxyl group.</u>

**Figure 6:**
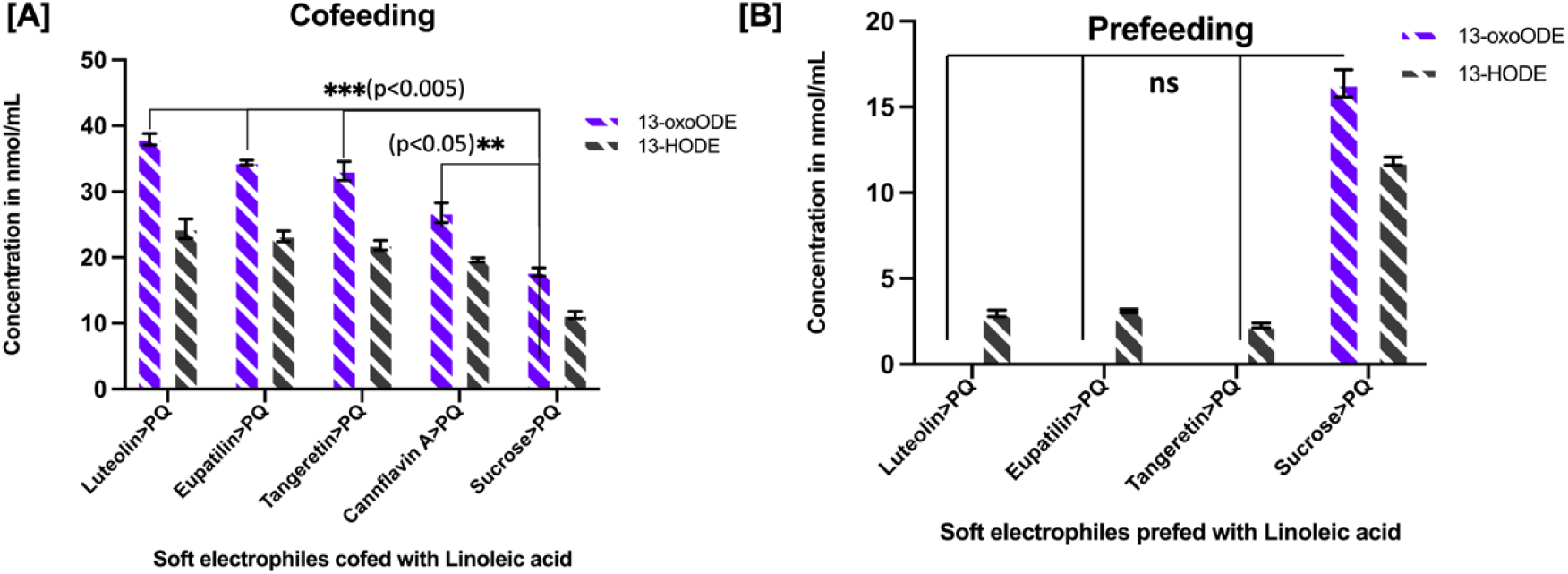
Differential feeding method in wild-type Canton S flies regulated flavone-mediated production of anti-inflammatory oxylipins post-PQ exposure: 200 *wild-type Canton S* adult male flies per feeding condition, aged 5 days post-emergence and differentially treated with a specific group of flavones, including Luteolin, Eupatilin, Tangeretin and Cannflavin, which have a similar chemical structure: **[A]** <u>There was *significant upregulation* of 13-oxoODE in the cofeeding method with Luteolin, Eupatilin and Tangeretin.</u> The concentration of 13-oxoODE was determined to be ∼48.446 nmol/mL in Luteolin-fed *wild-type Canton S* male fly heads, which is four-times the production as compared to 2.5% sucrose-fed fly heads (control). The data represents at least three biological replicates with at least two identical trials (n=200; Mean±SEM, \*\*\**p<0.005,**p<0.05* for one-way ANOVA). **[B]** <u>13-oxoODE was not produced in the prefeeding method, and 13-HODE production is much lower than in the 2.5% sucrose-fed fly heads (control) post-PQ exposure.</u>

Targeted lipidomic analysis using our specifically developed analytical method showed no significant upregulation of the linoleic acid–derived oxylipin 13-oxoODE compared with the control group. Similarly, none of the tested flavonoids significantly increased 13-HODE levels despite their structural. Consistent with these observations, our previous work demonstrated that flavonoids within this group fail to protect against PQ-induced phenotypes such as reduced lifespan, impaired mobility, elevated oxidative stress, and increased relish transcript expression [41].

Taken together, these findings suggest that additional structural determinants or specific molecular configurations are required to trigger anti-inflammatory responses in this model. The presence of individual electrophilic or hydroxyl functional groups alone appears insufficient to induce oxylipin production in response to toxin-induced inflammation.

### 2.5 Differential feeding method in *wild type Canton S* flies regulate flavone-mediated production of pro-resolving oxylipins post-PQ exposure

Two distinct feeding strategies were employed in this study because our previous work demonstrated that three specific flavones: Luteolin, Eupatilin, and Tangeretin did not exhibit protective effects when administered using the pre-feeding method alone [41]. In the pre-feeding approach, Canton S adult male flies were maintained for five days on standard fly food supplemented either with the respective flavones (Luteolin, Eupatilin, or Tangeretin; 1 mM) or with control diet, together with 1 mM linoleic acid. Following this pre-feeding period, flies were continuously exposed to 5 mM paraquat (PQ), and survival percentages were recorded every 24 hours until all flies had died.

In contrast, in the co-feeding method the flies were simultaneously exposed to the flavones (1 mM), linoleic acid (1 mM), and 5 mM PQ incorporated into the standard fly food, and survival was monitored at 24-hour intervals. Our previous study showed that the co-feeding regimen with these flavones significantly increased lifespan, improved locomotor performance, reduced oxidative stress, and suppressed relish transcript expression following PQ exposure [41]. Based on these findings, we applied both feeding paradigms to evaluate whether the timing of dietary supplementation influences oxylipin production following PQ-induced stress.

Targeted lipidomic analysis revealed that the pre-feeding method with Luteolin, Eupatilin, or Tangeretin did not result in detectable production or significant upregulation of the oxylipins 13-HODE and 13-oxoODE. In contrast, co-feeding these flavones together with linoleic acid during PQ exposure resulted in markedly elevated oxylipin levels compared with pre-feeding alone. These findings suggest that temporal synchronization between dietary intake and oxidative stress exposure is critical for optimal induction of oxylipin production, consistent with our previous observations involving polymethoxyflavones [41].

Interestingly, Cannflavin A exhibited a distinct pattern. Significant upregulation of 13-oxoODE was observed under both feeding conditions, with approximately 38.12 nmol/mL produced during pre-feeding and 27.58 nmol/mL during co-feeding in both adult male and female flies. This suggests that Cannflavin A may act through a mechanism that is less dependent on feeding timing, stress synchronization, or dietary intake behavior. Consequently, Cannflavin A represents a promising flavonoid candidate for further investigation in toxin-induced PD models.

Sex-specific analyses showed no major differences between male and female wild-type Canton S flies. Female flies similarly exhibited no significant oxylipin induction under the pre-feeding regimen with Luteolin, Eupatilin, or Tangeretin, whereas the co-feeding method produced significant increases in 13-oxoODE levels. In contrast, relish-null mutant flies of both sexes failed to show significant oxylipin upregulation following PQ exposure when the diet was supplemented with Luteolin, Eupatilin, Tangeretin, or Cannflavin A under the co-feeding regimen. Consistent with these findings, our previous work reported no significant improvement in lifespan or climbing ability in relish-null mutant males treated with these flavones [41], and similar observations were obtained in relish-null mutant females in the present study. Because these responses mirrored those observed in males, the corresponding female data are not presented separately.

### 2.6 Females show increased lifespan and produce more of the anti-inflammatory oxylipins in response to dietary supplementation with plant-derived soft electrophiles post-PQ exposure

Adult female *Canton S* flies showed a better survival rate compared to the control group, about 30% more than male flies at the interval of 48 hours post-PQ exposure. We also performed similar survival assays with *Canton S* adult female flies by supplementing the diet with Thymoquinone, Apigenin, DATQ, and Nobiletin in combination with Linoleic acid, which also showed an almost 30% increase in the survival rate of the female flies compared to their male counterparts (data not shown in the figure). On the contrary, Gardenin B, along with linoleic acid pretreatment, was detrimental to the survival percentage of both the adult male and female flies post-PQ exposure in comparison to the flies kept in control food with only Linoleic acid. Our previous findings supported this inference that Gardenin B, a structurally similar polymethoxyflavonoid, does not protect against toxicity and mobility impairments caused by PQ in *Canton S* adult male flies [36] and downregulates some major lipid classes in the cortical tissues of A53TSyn overexpressing mice [67]. Similar results were obtained when *Canton S* adult female flies were kept on a diet supplemented with compounds like Quercetin, Myrecetin, Fisetin, Hesperetin, and Naringin, which did not show any protective effects against PQ-induced toxicity and mobility impairment. This was previously shown as well in case of adult male flies [41] and thus is proved by the representative survival curve shown in **Fig. 7B**.

**Figure 7:**
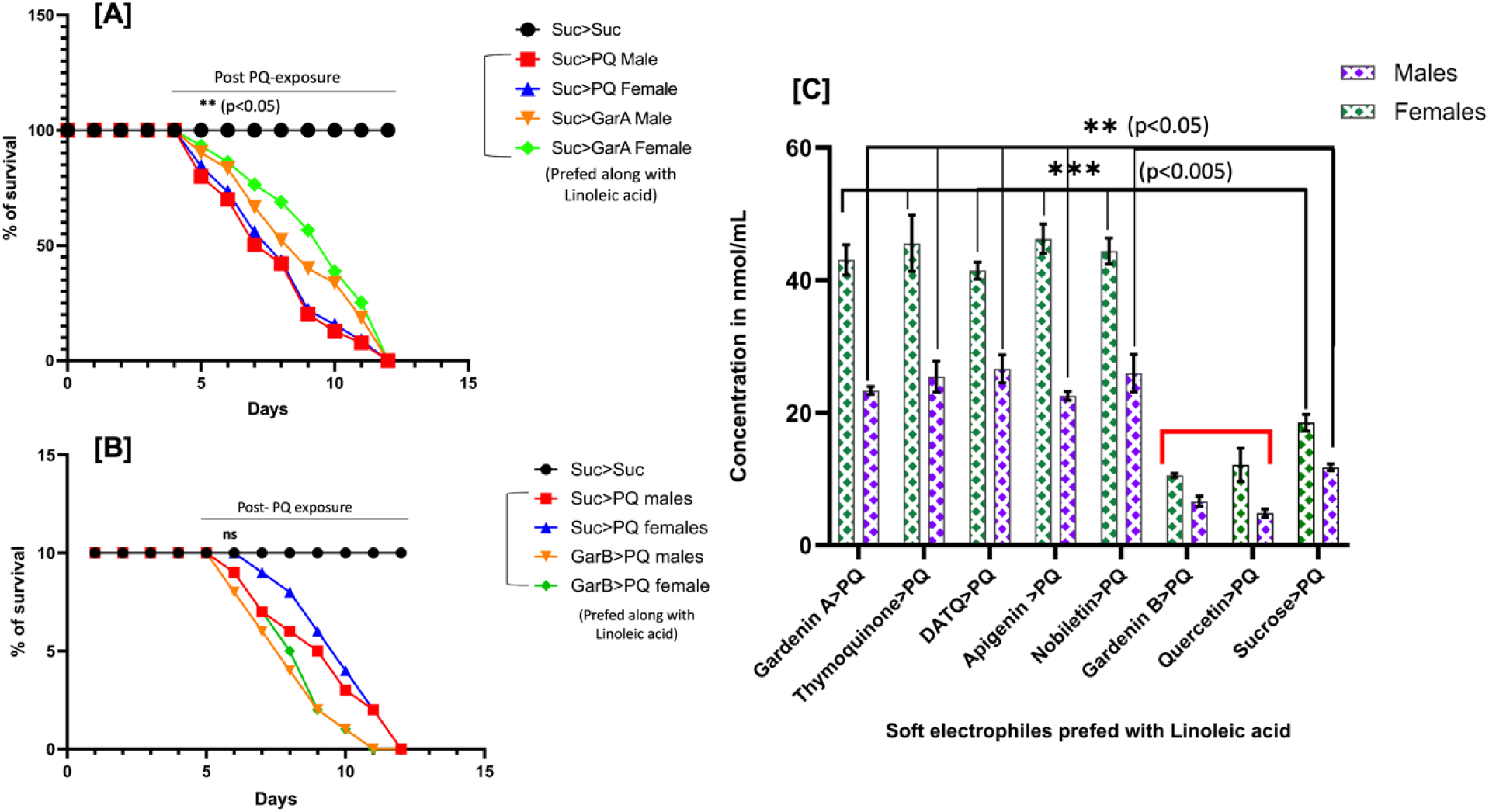
Females show increased lifespan and produce more of the pro-resolving oxylipins in response to dietary supplementation with plant-derived soft electrophiles post-PQ exposure: **[A]** <u>The graph represents the survival curve for adult males versus females kept on a diet supplemented with Gardenin A and linoleic acid.</u> 10 separate biological replicates with 10 flies per feeding condition are shown in the data. The data displayed is mean ± SEM. *p < 0.05 according to the Kaplan-Meier survival test. **[B]** <u>The graph represents the survival curve for adult males versus females kept on a diet supplemented with Gardenin B and Linoleic acid.</u> 10 separate biological replicates with 10 flies per feeding condition are shown in the data. The data displayed is mean ± SEM. Gardenin B *did not significantly* improve the lifespan of both male and female flies post-PQ exposure. **[C]** <u>The graph represents significantly higher upregulation of 13-oxoODE and 13-HODE in *wild-type Canton S* adult female fly heads (n=200) as compared to adult male fly heads.</u> The average concentration of 13-oxoODE and13-HODE was ∼41.221 nmol/mL in female fly heads and 23.308 nmol/mL in adult male fly heads as compared to 2.5% Sucrose (control group) post-PQ exposure. There was *no significant upregulation* in the level of oxylipins following dietary supplementation with Gardenin B and Quercetin (indicated in **red bar**). The data represents at least three biological replicates with at least two identical trials (Mean ± SEM, **p<0.05, *\*\*\**p<0.005 for one-way ANOVA).

A significant upregulation in the levels of both 13-HODE and 13-oxoODE has been observed in the adult female flies post-PQ exposure as compared to the adult male flies. The average concentration of 13-oxoODE+13-HODE was found to be 41.221 nmol/mL (3.26 ng/head) for Gardenin A-fed female fly heads as compared to 23.308 nmol/mL (1.61ng/head) in the adult male fly heads post-PQ exposure, which indicated almost **double the upregulation of oxylipins** in general in the female fly heads as shown in **Fig. 7C**. In contrast, compounds like Gardenin B, Quercetin, Myrecetin, Fisetin, Hesperetin, and Naringin failed to further upregulate the average concentration of oxylipins in the males and females unanimously. A representative quantity for all these compounds having a saturated C-ring in their chemical structure is indicated by the **red bar** in **Fig. 7C**. To model the higher incidence of PD observed in human males, most of the previous studies regarding PD and its effects have successfully utilized adult male flies. Male flies have been reported to exhibit greater sensitivity to PQ-induced toxicity and develop PD-like phenotypes earlier and more robustly than female flies, including locomotor deficits and neurodegenerative changes [36,41,65]. Therefore, the use of male flies provides a biologically relevant system for examining PQ-induced neurotoxicity and related pathological processes. This sex-specific sensitivity is in line with mammalian model studies showing that male bias in experimental PD results from both male-specific genetic variables and the neuroprotective effects of estrogen in females [78]. Together, these findings indicate that both sex hormones and sex chromosome–encoded factors contribute to sexual dimorphism in PD vulnerability, supporting our observation of considerably higher oxylipin upregulation in female flies as compared to males. This also posits the extensive use of male counterparts as the more biologically relevant model for investigating PQ-induced neurotoxicity.

## Discussion

Recent research increasingly highlights the roles of environmental exposures, gut microbiota composition, and genetic susceptibility in the onset and progression of Parkinson’s disease (PD), particularly in aging populations [6–9]. A common consequence of these interacting factors is chronic neuroinflammation, now widely recognized as a central pathological hallmark of PD. Sustained activation of innate immune glial cells, together with elevated levels of inflammatory mediators in brain tissue, perpetuates a self-reinforcing inflammatory environment that promotes neuronal dysfunction and degeneration, ultimately leading to PD-like phenotypes [45]. Importantly, lifestyle interventions, including regular physical activity and diets enriched in natural bioactive compounds have been shown in multiple preclinical models to attenuate neuroinflammation and reduce PD risk [10–13]. These observations emphasize the importance of promoting active resolution of neuroinflammation as a potential preventive and disease-modifying strategy, particularly in aging-associated neurodegeneration.

Resolution of inflammation is an active and tightly regulated biological process involving complex molecular signaling networks. In mammals, this process is mediated in part by specialized pro-resolving lipid mediators (SPMs) derived from essential omega-3 and omega-6 fatty acids (FAs), including lipoxins, resolvins, maresins, and protectins. These lipid mediators orchestrate the termination of inflammatory responses by promoting tissue repair, restoring homeostasis, and limiting excessive immune activation [46–48]. Importantly, lipid-derived pro-resolving oxylipins are not restricted to mammals; analogous molecules have also been identified in model organisms such as *Drosophila melanogaster* [30,31,38]. Like mammals, flies cannot synthesize essential FAs de novo and must obtain them through the diet [31,38]. Evidence from diverse model systems and clinical studies therefore supports the concept that inflammation resolution represents an evolutionarily conserved process mediated by lipophilic signaling molecules derived from essential FAs, as well as by dietary lipophilic electrophiles whose metabolic and signaling functions appear conserved across species [49,50].

Our findings further highlight the role of plant-derived lipophilic soft electrophiles in enhancing the pro-resolving capacity of linoleic acid-derived oxylipins in *Drosophila*. Mechanistically, the structural properties of these compounds, particularly their lipophilicity and electrophilic moieties likely facilitate interactions with cellular membranes and influence oxylipin biosynthetic pathways [51]. Notably, despite the abundance of omega-3 and omega-6 FA precursors within cellular membranes, reduced oxylipin levels have been reported in both PD patients and experimental PD models [52]. Such deficiencies may arise from impaired FA metabolism, altered lipid mediator balance favoring pro-inflammatory pathways, or limited substrate availability [53]. Consequently, growing attention has focused on therapeutic strategies aimed at enhancing the resolution phase of inflammation through increased production of pro-resolving oxylipins [54–57]. Consistent with this concept, epidemiological studies indicate that higher dietary intake of omega-3 and certain omega-6 polyunsaturated FAs is associated with a reduced risk of neurodegenerative diseases [23,26,27].

Building on this framework, we hypothesized that regular dietary intake of polymethoxyflavones and related plant-derived soft electrophiles could enhance oxylipin biosynthesis and thereby limit chronic neuroinflammation. To test this hypothesis, we developed and validated a sensitive analytical method tailored for *Drosophila*, enabling accurate quantification of low-abundance lipid-derived mediators in neural tissue. Using this approach, we show that dietary supplementation with soft electrophiles significantly increased levels of the pro-resolving oxylipin 13-oxoODE, particularly in the presence of linoleic acid as a metabolic precursor. These findings indicate that oxylipin production in flies is regulated by both substrate availability and electrophile-triggered signaling mechanisms.

Comparative analyses of oxylipin pathways across biological kingdoms reveal remarkable evolutionary conservation between plants and animals [58], suggesting that certain plant secondary metabolites may function as interspecies chemical signals involved in stress-response pathways. Although natural products often display diverse biological activities, elucidating their precise mechanisms of action remains challenging compared with synthetic compounds. Flavonoids and related phytochemicals have long been investigated for their antioxidant and anti-inflammatory properties [59,60], with early studies focusing on their capacity to reduce oxidative stress in neurodegenerative models [61–63]. However, many flavonoids exhibit pleiotropic reactivity and frequently produce false-positive results in in vitro assays, leading to their classification as PAINS (pan-assay interference compounds) and limiting their development as conventional drug candidates [64].

Nevertheless, increasing in vivo and clinical evidence suggests that the beneficial effects of flavonoids are not primarily mediated by direct antioxidant activity [65]. In models of paraquat-induced oxidative stress, antioxidant capacity alone appears insufficient to confer neuroprotection [6,36]. Instead, flavonoids are increasingly recognized as modulators of conserved cellular signaling pathways and lipid metabolic networks rather than classical radical scavengers [66,67]. From an evolutionary perspective, many plant polyphenols likely evolved as defense molecules against herbivores and environmental stress and were subsequently co-opted in animals as signaling modulators. Plant-derived salicylates provide a notable example. Aspirin (acetylsalicylic acid), originally isolated from *Salix alba*, not only inhibits cyclooxygenase-mediated prostaglandin synthesis but also promotes the formation of electrophilic pro-resolving lipid mediators (EFOX) and facilitates inflammation resolution [68,69].

Consistent with this concept, aspirin has been shown in *Drosophila* to regulate COX-like activity and stimulate the production of the linoleic acid–derived oxylipins 13-HODE and 13-oxoODE, thereby linking NF-κB signaling to eicosanoid metabolism [31]. In line with these observations, our findings show that structurally related plant-derived soft electrophiles also enhance oxylipin biosynthesis and promote resolution signaling in a paraquat-induced model of neuroinflammation. Complementary studies further show that flavonoids such as xanthohumol and isoquercetin regulate lipid droplet biogenesis in *Drosophila*, linking phytochemicals to cellular stress adaptation and inflammatory control [51].

Although polyphenols rarely conform to classical drug-likeness criteria, historical and biological evidence suggests that certain plant-derived compounds, such as vitamins are essential for maintaining physiological homeostasis [70,71]. Szent-Györgyi’s concept of “vitamin P,” for example, emphasized the importance of plant secondary metabolites in supporting vitamin function [64]. Consistent with this view, structurally related compounds such as curcumins, natural chalcones, and vitamin E derivatives have been shown to influence lipid oxidation pathways and promote resolution of inflammation [72–74].

In the present study, we therefore focused on a defined subset of plant-derived soft electrophiles containing α,β-unsaturated carbonyl groups and evaluated their effects as dietary supplements. Compounds examined included Gardenin A, Thymoquinone, d-α-Tocopherylquinone, Puresirtmax, and Cannflavin A, each characterized by electrophilic and lipophilic structural features associated with anti-inflammatory activity. All tested electrophiles significantly improved lifespan, rescued paraquat-induced locomotor deficits, and markedly increased oxylipin production. Notably, co-supplementation with linoleic acid resulted in an approximately four-fold increase in oxylipin levels compared with linoleic acid alone, suggesting that these compounds function as metabolic cofactors that enhance pro-resolving lipid biosynthesis.

Importantly, linoleic acid supplementation in the absence of soft electrophiles favored alternative oxidative pathways, including formation of malondialdehyde, a reactive aldehyde known to exacerbate neuroinflammation [75]. In contrast, the presence of electrophiles redirected metabolic flux toward bioactive pro-resolving oxylipins and promoted lipid mediator class switching, thereby amplifying resolution capacity [76]. These findings underscore the critical interplay between dietary composition, molecular structure, and lipid-mediated inflammation resolution.

Structure–activity analyses further demonstrated that electrophilic reactivity is essential for bioactivity *in vivo*. Compounds lacking the α,β-unsaturated carbonyl moiety or containing substitutions at the C3 position failed to induce oxylipin production, consistent with previous reports [41]. This observation contrasts with traditional *in vitro* antioxidant assays, which emphasize hydroxylation patterns as determinants of activity, highlighting the limitations of reductive antioxidant paradigms for predicting in vivo efficacy.

At the molecular level, our results demonstrate that oxylipin induction requires the NF-κB homolog Relish. Relish-null mutants failed to produce 13-HODE and 13-oxoODE and exhibited no improvement in lifespan or locomotor performance despite dietary supplementation, revealing a link between innate immune signaling, redox biology, and lipid metabolism. These findings are consistent with extensive evidence implicating NF-κB pathways in aging, neuroinflammation, and neurodegeneration [6,36,67]. Interestingly, sex-specific differences were also observed, with female flies displaying higher oxylipin levels and greater longevity benefits than males. This observation parallels known sex-dependent immune and metabolic responses and reflects epidemiological data indicating differences in PD incidence and progression between men and women [77,78]. Hormonal regulation, metabolic differences, and gut microbiota composition may contribute to these effects [79,80].

Finally, the paraquat-induced *Drosophila* model employed in this study provided an effective platform for integrating phenotypic, molecular, and analytical lipidomics approaches. Given the strong conservation of inflammatory and metabolic pathways between flies and humans, and the inability of *Drosophila* to synthesize essential polyunsaturated fatty acids de novo, this system offers a powerful model for investigating diet–gene–environment interactions in neuroinflammation. The development of a sensitive LC–MS-based method for oxylipin quantification in fly heads also addresses long-standing analytical challenges related to the low abundance of these lipid mediators.

Collectively, our findings support the existence of a conserved and inducible oxylipin-mediated resolution system modulated by diet, genetics, and oxidative stress. These results provide new mechanistic insight into how plant-derived soft electrophiles regulate lipid-mediated inflammation resolution and highlight their potential as dietary modulators of neuroinflammatory processes. Such insights may ultimately inform nutritional and metabolic strategies aimed at preventing or delaying neurodegenerative diseases.

## Materials and Methods

### Drosophila Culturing and Stocks

The fly stocks were cultured on a standard medium based on the recipe from the Bloomington Stock Center. This medium included ingredients like cornmeal, corn syrup, yeast, and agar. The rearing took place at a temperature of 25 °C under a cycle of 12 hours of light followed by 12 hours of darkness. The control genotype for all experiments was the wild-type strain known as *Canton S*. Male and female flies aged between 3 to 5 days post-emergence were selected for all the tests. Some of the stocks used in the study, namely *CantonS* and *RelE20*, were obtained from the Bloomington Drosophila Stock Center at Indiana University.

### Chemicals

Various chemicals were procured from Sigma-Aldrich, including Sucrose, Paraquat (methyl viologen dichloride hydrate), Linoleic acid, Gardenin A (GA), Gardenin B (GB), Thymoquinone (TQ), D-alpha-tocopherylquinone (DQ), Apigenin (AP), Nobiletin (NB), Puresirtmax (PX), Cannflavin A (CA), Quercetin (QR), Myrecetin (MYR), Fisetin (FS), Hesperitin (HS), Naringin (NR), Luteolin (LT), Eupatilin (EP) and Tangeretin (TG). Additionally, Cayman chemicals provided 13-HODE and 13-oxoODE along with the internal standard, ^13^C-labelled Linoleic acid.

### Other materials required

Solid phase extraction STRATA (SPE) cartridges (Phenomenex), Non-chiral reverse phase HPLC columns, MS grade: water, acetonitrile (50:50 acidified with 0.5% TFA), and formic acid; Analytical grade standards: 13-HODE, 13-oxoODE and ^13^C-labelled Linoleic acid (Cayman chemicals), Other chemicals: methanol, hexane, ethanol, methyl formate, dimethyl sulfoxide, sucrose, paraquat (methyl viologen) etc.

### Fly Food Preparation

A solid food mixture was prepared by combining 175.67g of Bloomington Nutrifly powder with 1 liter of distilled water. Tegosept stock solution (10mL) and propionic acid (4.8mL) were added to the mixture. 5mL of the solid food mixture was dispensed into each feeding vial. 20 male/female adult flies (Canton S or *relish* null) were transferred into each vial. A stock solution of 10mM concentration is prepared in 2.5% sucrose with 10mM of Linoleic acid and 10mM of each sample solution. A final concentration of 1mM of the (Linoleic acid+test sample/2.5% sucrose) solution is maintained in the fly food in each vial. So, a 500 μL aliquot of each sample solution is added to the freshly prepared fly food in each vial while the food is still moderately hot and mixed well. The following samples were used for the dietary supplementation in the fly food from a stock of 100mM in DMSO and diluted to 10mM with 2.5% sucrose: Linoleic acid (LA), Gardenin A (GA), Gardenin B (GB), Thymoquinone (TQ), D-alpha-tocopherylquinone (DQ), Apigenin (AP), Nobiletin (NB), Puresirtmax (PX), Cannflavin A (CA), Quercetin (QR), Myrecetin (MYR), Fisetin (FS), Hesperitin (HS), Naringin (NR), Luteolin (LT), Eupatilin (EP) and Tangeretin (TG).

### Drosophila Feeding Regimen, Survival and mobility assays

Adult male/female flies (5 days post-eclosion) were collected and maintained in standard fly vials.10 flies per vial were maintained in case of survival and mobility assays. For feeding, the standard fly diet was supplemented with sample solutions containing a mixture of linoleic acid (LA) and either specific plant-derived soft electrophiles or 2.5% sucrose of equal volume (used as the control) at an overall 1mM concentration. Flies were kept in the supplemented diet for 5 days, and mortality was recorded from the initiation of PQ exposure (Day 0) under both pre-feeding and co-feeding paradigms. For paraquat exposure in pre-feeding condition, flies were transferred onto new vials prepared using filter paper saturated with 5mM Paraquat (methyl viologen) after 5 days. These vials were changed every alternate day, and all the dying flies were discarded. For the cofeeding condition, 5mM paraquat was mixed with the aliquot solution of 1mM of LA and specific samples or 2.5% sucrose together in the standard fly diet. Survival was monitored daily until all flies in each condition had died. A new batch of food was prepared every alternate day to make sure that the dead flies are being discarded at regular intervals. Each experimental condition was replicated in at least five independent biological replicates, each consisting of 10 flies.

Negative geotaxis assays were conducted to assess locomotor deficits in adult male/female flies subjected to different feeding conditions. For each condition, 10 flies were transferred into an empty plastic vial and gently tapped to the bottom. The number of flies that climbed past a 5 cm mark within 30 seconds was recorded and expressed as a percentage of the total. Climbing performance was recorded 24 hours and 48 hours after paraquat (PQ) exposure under both pre-feeding and co-feeding paradigms. Each condition was tested using five independent biological replicates, with each replicate measured in triplicate at 5-minute intervals. The average climbing percentages were calculated and plotted. Statistical analysis was performed using one-way ANOVA, with significance set at p < 0.05.

### Separation of Fly Heads

For separation of fly heads, whole flies were first transferred to Eppendorf tubes (50 flies per tube). CO_2_-anethesized flies in the Eppendorf tubes were kept in liquid nitrogen for about 1-2 mins for flash freezing the organisms. The tubes were shaken vigorously inside the liquid nitrogen container and then tapped on the bench top so that individual body parts separate out. The broken flies were transferred onto a fine sieve mesh from where the isolated heads were carefully collected with forceps. The fly samples were processed in groups of 100 heads per Eppendorf tube for further analysis.

### Quantitative real time PCR

To assess gene expression of *relish*, total RNA was extracted from the heads of 30 adult male flies using TRIzol reagent (Invitrogen), following standard protocols from the manufacturer. cDNA synthesis was carried out using 1 µg of RNA as input with the High Capacity cDNA Reverse Transcription Kit (Applied Biosystems) as defined in the manufacturer’s protocols. Quantitative rt-PCR was conducted using SYBR supermix (iQ SYBR Green Supermix, Bio-Rad) on a StepOnePlus Real-Time PCR System. Transcript abundance was determined by the comparative Ct method (ΔΔCt), with ribosomal protein L32 (*RpL32/RP49*) serving as the internal reference gene. For each experimental feeding condition, gene expression analysis was performed using at least three independent biological replicates. The following are the primer sequences used: rp49 (F): 5′ AGA AGC GCA AGG AGA TTG TC 3′; rp49(R): 5′ ATG GTG CTG CTA TCC CAA TC 3′; relish(F): 5′ GGC ATC ATA CAC ACC GCC AAG AAG 3; relish(R): 5′ GTA GCT GTT TGT GGG ACA ACT CGC 37

### Extraction of Lipids

For the extraction of the lipids and subsequent QTOF analysis, 10 vials with 20 flies in each were considered for each sample or control batch, resulting in a total of 200 flies per feeding condition. 200 fly head samples for each category were first washed with DI water, 70% ethanol, and again with DI water. Samples were then air dried and transferred to new pre-chilled Eppendorf tubes (100 head samples per tube). Samples were then ground and homogenized at 0°C with the pestle in 20 µL of 15% cold methanol to extract the lipid content in methanol. The homogenate was left on ice in dark for 15 mins. 10 μL of the 10 µM stock solution of internal standard (^13^C-linoleic acid) was added to 20 μL of the homogenate to achieve an overall concentration of 5 μM internal standard. The homogenate was then centrifuged at 4°C and 3700g for 15 minutes. Supernatant was finally recovered and kept on ice in dark for 15 mins. The pH was adjusted to 3.0 by adding 10-20 drops of dilute HCl. SPE columns were first conditioned with a mixture of 20 mL water + 20 mL methanol. The columns were then washed with a mixture of 20 mL ice cold 15%(v/v) methanol + 20 mL water + 20 mL n-hexane. The recovered supernatant was immediately loaded onto the SPE column. Lipids were recovered on 500 μL of cold methyl formate solution (80% v/v). Excess methyl formate was evaporated using a rotatory evaporator. Lipids were dissolved in 500 µL of cold degassed methanol and can be stored at -80 °C for further use (better to use the freshly prepared extract). For each experimental feeding condition, QTOF analysis was performed using at least three independent biological replicates and two technical replicates.

### Preparation of Calibration Standards

Matrix-matched calibration standards were prepared using the isotopic dilution method with different concentrations of 13-HODE and 13-oxoODE separately (10, 7.5, 5.0, 2.5, 1.25, 0.625, 0.3125 µg/mL, equivalent to 33.73, 25.30, 16.87, 8.43, 4.22, 2.11, 1.05 nmol/mL for 13-HODE and 33.96, 25.47, 16.98, 8.49, 4.25, 2.12, 1.06 nmol/mL for 13-oxoODE) in the fly head homogenate. This homogenate was extracted from control *wild-type Canton S* male fly heads kept in standard fly food for 7 days. Internal standard, a ^13^C-labelled linoleic acid (Cayman Chemicals), was added to the standard solutions at a final concentration of 5 μM, all prepared in triplicates. A separate set of calibration standards in the same concentration range and 5 μM final concentration of internal standard was prepared in methanol. The matrix effect was calculated as a percentage ratio of the slope of the fly homogenate standard curve and methanol standard curve to account for the matrix effect of fly head homogenate. All calibration sets were processed in triplicate and two technical repeats.

### Analysis and quantification of Lipids in UPLC-QTOF-HRMS

Lipid UPLC-HRMS analysis was performed using a Waters Xevo G2-XS QTOF mass spectrometer connected to an ACUITY I-Class UPLC system. An ACUITY UPLC BEH C18 column (2.1 x 50 mm, 1.7 mm) was used with an ACUITY UPLC BEH C18 Van Guard Pre-column (2.1 x 5 mm, 1.7 mm). The column was eluted at flow rate of 0.3 mL/min with mobile phases A (water with 0.1% formic acid) and B (acetonitrile with 0.1% formic acid) gradient: 35% B for 1 minute, changing from 35% to 90% B over 10 minutes, shifting from 90% to 100% B for 0.1 minutes, holding at 100% B for 2 minutes, returning from 100% to 35% B for 0.1 minutes, and finally holding at 35% B for 1.2 minutes. The total run time for each sample was maintained at 14 minutes. The ESI source conditions included: Capillary voltage 2.15V, Sampling cone 40 V, Source offset 80, Source temperature 10 °C, Desolvation temperature 36 °C, Cone gas 15 L/h, Desolvation gas 400 L/h. The lock mass at 554.2615 in negative resolution mode (leucine enkephalin, peak resolution 25,000). The column temperature was 35°C, and sample temperature was 10°C.

Matrix-matched and in-solvent calibration curves were used for quantification of 13-HODE and 13-oxoODE. Extracted ion chromatograms within 10ppm error window were obtained for 13-HODE and 13-oxoODE and quantified relative to the isotopically labeled internal standard (^13^C-LA). Peak area ratios of analytes to internal standard were plotted against concentrations of 13-HODE/13-oxoODE to calculate the relative quantities of oxylipins. Limits of detection (LOD) and quantification (LOQ) were calculated in accordance with International Conference on Harmonisation (ICH) guidelines (refer to **Appendix A Fig.2** for the verification of calculation). Quantitative measurements were derived from pooled biological samples with three biological replicates and two technical replicates. 13-HODE was detected at 295.2273 m/z, 13-oxoODE at 293.2117 m/z and internal standard (^13^C-LA) at 297.2928 m/z. The relative quantities of both 13-HODE and 13-oxoODE were calculated using the formula: [43 nmol/mL (derived concentration) * 0.05 mL (volume of homogenate used) * 293.2 (MW)] / 200 (no. of fly heads) = 3.15 ng/head (for 13-oxoODE). Percent recovery was calculated using the following equation: % Recovery = ((Observed Concentration – Endogenous Concentration) / Spiked Diluent Concentration) × 100. Here, the observed concentration refers to the analyte concentration measured in the spiked sample, the endogenous concentration corresponds to the analyte level in the unspiked sample, and the spiked diluent concentration denotes the known quantity of analyte added [44].

### Statistics and software

GraphPad Prism 11 (GraphPad Software, Inc., La Jolla, CA, USA) was used for all statistical analyses. The data are displayed as mean ± standard error of the mean (SEM), and SEM error bars are incorporated in all the linear as well as bar graphs. The survival assays Depending on the experimental design, either a one-way or two-way analysis of variance (ANOVA) was used to evaluate statistical significance between experimental groups. The survival assays were analysed using the Kaplan-Meier non-parametric test. At p<0.5, effects were determined to be statistically significant. The following notation is used in the figures to highlight statistically significant differences: *p<0.5, **p<0.05, ***p<0.005. Relative standard deviation (RSD) studies were performed for analytical measurements in order to assess the accuracy and repeatability of quantified data. Matrix-matched calibration curves produced by linear regression analysis were used to quantify 13-HODE and 13-oxoODE in order to identify correlation coefficients and best-fit standard equations. The graphical abstract was prepared in BioRender.

## Conclusion and Future works

In the context of innate immunity response pathways, a very significant aspect is the interaction of lipophilic soft electrophiles with cellular components. These interactions primarily involve electrophile-responsive regulatory proteins and membrane lipids that are rich in nucleophilic regions, often composed of cysteine thiols. A noteworthy feature is the structural resemblance of these lipophilic dietary molecules to the oxylipins, which might facilitate the resolution of neuroinflammation. In summary, the converging structural attributes and consistent functional outcomes underscore the potential therapeutic implications of lipophilic soft electrophiles in addressing conditions related to neuroinflammation. Nevertheless, the majority of the information now available comes from experimental models, which are unable to accurately capture patient variability, age-related metabolic changes, or the complexity of the human brain. Therefore, extensive validation in physiologically relevant models, long-term human trials, and meticulous assessment of safety, dosage, and delivery will be necessary to translate these findings into clinical practice. However, further research is being undertaken to visualize the localization and spatial distribution of these essential lipophilic soft electrophiles using highly advanced mass spectrometric techniques, to better understand their mechanistic actions and develop potential preventative and/or curative strategies for PD.

## Acknowledgement

This work was partially supported by the Office of Dietary Supplements and National Center for Complimentary and Integrative Medicine of the National Institutes of Health under awards number 1R41AT011716–01 (Ciesla) and R03AT011871–01 (Ciesla & Maitra).

## Author contributions

Conceptualization – Swarnali Chatterjee, Urmila Maitra, and Lukasz Ciesla; data acquisition –Swarnali Chatterjee, Bianca McCarty, Caleb Vandenberg, Madison Bever; data analysis and visualization – Swarnali Chatterjee, Qiaoli Liang, Lukasz Ciesla; manuscript preparation – Swarnali Chatterjee, Qiaoli Liang, Urmila Maitra, Lukasz Ciesla; funding acquisition – Lukasz Ciesla, Urmila Maitra.

**Appendix A**. Supplemental information

**Appendix B.** Figures in high resolution

## Data availability

Data will be made available upon request from the corresponding author.

## References

1. Dorsey, E. R., Sherer, T., Okun, M. S., & Bloem, B. R. (2018). The emerging evidence of the Parkinson pandemic. Journal of Parkinson’s Disease, 8(s1), S3–S8.

2. Yang, W., Hamilton, J. L., Kopil, C., Beck, J. C., Tanner, C. M., Albin, R. L., Ray Dorsey, E., Dahodwala, N., Cintina, I., & Hogan, P. (2020). Current and projected future economic burden of Parkinson’s disease in the US. Npj Parkinson’s Disease, 6(1), 15.

3. Breydo, L., Wu, J. W., & Uversky, V. N. (2012). α-Synuclein misfolding and Parkinson’s disease. Biochimica et Biophysica Acta (BBA)-Molecular Basis of Disease, 1822(2), 261–285.

4. Soares, J. J., Rodrigues, D. T., Gonçalves, M. B., Lemos, M. C., Gallarreta, M. S., Bianchini, M. C., Gayer, M. C., Puntel, R. L., Roehrs, R., & Denardin, E. L. (2017). Paraquat exposure-induced Parkinson’s disease-like symptoms and oxidative stress in Drosophila melanogaster: Neuroprotective effect of Bougainvillea glabra Choisy. Biomedicine & Pharmacotherapy, 95, 245–251.

5. Tong, T., Duan, W., Xu, Y., Hong, H., Xu, J., Fu, G., Wang, X., Yang, L., Deng, P., & Zhang, J. (2022). Paraquat exposure induces Parkinsonism by altering lipid profile and evoking neuroinflammation in the midbrain. Environment International, 169, 107512.

6. Maitra, U., Scaglione, M. N., Chtarbanova, S. & O’Donnell, J. M. (2019) Innate immune responses to paraquat exposure in a Drosophila model of Parkinson’s disease. Scientific Reports 9, 12714.

7. Zaman, V., Shields, D. C., Shams, R., Drasites, K. P., Matzelle, D., Haque, A., & Banik, N. L. (2021). Cellular and molecular pathophysiology in the progression of Parkinson’s disease. Metabolic brain disease, 36, 815–827.

8. Zhang, W., Phillips, K., Wielgus, A. R., Liu, J., Albertini, A., Zucca, F. A., Faust, R., Qian, S. Y., Miller, D. S., Chignell, C. F., Wilson, B., Jackson-Lewis, V., Przedborski, S., Joset, D., Loike, J., Hong, J.-S., Sulzer, D., & Zecca, L. (2011). Neuromelanin Activates Microglia and Induces Degeneration of Dopaminergic Neurons: Implications for Progression of Parkinson’s Disease. Neurotoxicity Research, 19(1), 63–72.

9. Forman, M. S., Trojanowski, J. Q., & Lee, V. M. (2004). Neurodegenerative diseases: a decade of discoveries paves the way for therapeutic breakthroughs. Nature medicine, 10(10), 1055–1063.

10. Neshatdoust, S., Saunders, C., Castle, S. M., Vauzour, D., Williams, C., Butler, L., Lovegrove, J. A., & Spencer, J. P. E. (2016). High-flavonoid intake induces cognitive improvements linked to changes in serum brain-derived neurotrophic factor: Two randomised, controlled trials. Nutrition and Healthy Aging, 4(1), 81–93.

11. Neshatdoust, S., Saunders, C., Castle, S. M., Vauzour, D., Williams, C., Butler, L., Lovegrove, J. A., & Spencer, J. P. E. (2016). High-flavonoid intake induces cognitive improvements linked to changes in serum brain-derived neurotrophic factor: Two randomised, controlled trials. Nutrition and Healthy Aging, 4(1), 81–93.

12. Godos, J., Scazzina, F., Paternò Castello, C., Giampieri, F., Quiles, J. L., Briones Urbano, M., Battino, M., Galvano, F., Iacoviello, L., De Gaetano, G., Bonaccio, M., & Grosso, G. (2024). Underrated aspects of a true Mediterranean diet: Understanding traditional features for worldwide application of a “Planeterranean” diet. Journal of Translational Medicine, 22(1), 294.

13. Zu, Y., Zhang, X., & Feng, C. (2022). Effect of enalapril maleate-folic acid tablets on inflammatory response and myocardial endoplasmic reticulum stress-related factors in hypertensive rats. Tropical Journal of Pharmaceutical Research, 21(5), 1003–1008.

14. Dini, I., & Grumetto, L. (2022). Recent Advances in Natural Polyphenol Research. Molecules (Basel, Switzerland), 27(24), 8777.

15. Sergent, T., Piront, N., Meurice, J., Toussaint, O., & Schneider, Y. J. (2010). Anti-inflammatory effects of dietary phenolic compounds in an in vitro model of inflamed human intestinal epithelium. Chemico-Biological Interactions, 188(3), 659–667.

16. Fakhri, S., Piri, S., Moradi, S. Z., & Khan, H. (2022). Phytochemicals targeting oxidative stress, interconnected neuroinflammatory, and neuroapoptotic pathways following radiation. Current Neuropharmacology, 20(5), 836.

17. Jamieson, P. E., Smart, E. B., Bouranis, J. A., Choi, J., Danczak, R. E., Wong, C. P., Paraiso, I. L., Maier, C. S., Ho, E., Sharpton, T. J., Metz, T. O., Bradley, R., & Stevens, J. F. (2024). Gut enterotype-dependent modulation of gut microbiota and their metabolism in response to xanthohumol supplementation in healthy adults. Gut Microbes, 16(1), 2315633.

18. Campolo, M., Paterniti, I., Siracusa, R., Filippone, A., Esposito, E., & Cuzzocrea, S. (2019). TLR4 absence reduces neuroinflammation and inflammasome activation in Parkinson’s diseases in vivo model. Brain, behavior, and immunity, 76, 236–247.

19. Talevi A. (2015). Multi-target pharmacology: possibilities and limitations of the “skeleton key approach” from a medicinal chemist perspective. Frontiers in pharmacology, 6, 205.

20. Patel, J., Pramanik, A., Maheshwari, S., Acharya, N., & Pathak, Y. (2026). Safety and Toxicity Aspects for Nutraceuticals and Related Products: Regulatory Provisions across the Globe. In Nutraceuticals (pp. 54–81). CRC Press.

21. LoPachin, R. M., Geohagen, B. C., & Nordstroem, L. U. (2019). Mechanisms of soft and hard electrophile toxicities. Toxicology, 418, 62–69.

22. Zhang, J., Wang, C., Ji, L., & Liu, W. (2016). Modeling of toxicity-relevant electrophilic reactivity for guanine with epoxides: estimating the hard and soft acids and bases (HSAB) parameter as a predictor. Chemical Research in Toxicology, 29(5), 841–850.

23. Andrade, Y. M. F. de S., Castro, M. V. de, Tavares, V. de S., Souza, R. da S. O., Faccioli, L. H., Lima, J. B., Sorgi, C. A., Borges, V. M., & Araújo-Santos, T. (2023). Polyunsaturated fatty acids alter the formation of lipid droplets and eicosanoid production in Leishmania promastigotes. Memórias Do Instituto Oswaldo Cruz, 118.

24. Cipollina, C. (2015). Endogenous generation and signaling actions of omega-3 fatty acid electrophilic derivatives. BioMed research international, 2015.

25. Labrousse, V. F., Nadjar, A., Joffre, C., Costes, L., Aubert, A., Grégoire, S., Bretillon, L., & Layé, S. (2012). Short-Term Long Chain Omega3 Diet Protects from Neuroinflammatory Processes and Memory Impairment in Aged Mice. PLoS ONE, 7(5), e36861.

26. Hernando, S., Requejo, C., Herran, E., Ruiz-Ortega, J. A., Morera-Herreras, T., Lafuente, J. V., Ugedo, L., Gainza, E., Pedraz, J. L., Igartua, M., & Hernandez, R. M. (2019). Beneficial effects of n-3 polyunsaturated fatty acids administration in a partial lesion model of Parkinson’s disease: The role of glia and NRf2 regulation. Neurobiology of Disease, 121, 252–262.

27. Zendedel, A., Habib, P., Dang, J., Lammerding, L., Hoffmann, S., Beyer, C., & Slowik, A. (2015). Omega-3 polyunsaturated fatty acids ameliorate neuroinflammation and mitigate ischemic stroke damage through interactions with astrocytes and microglia. Journal of neuroimmunology, 278, 200–211.

28. Abrescia, P., Treppiccione, L., Rossi, M., & Bergamo, P. (2020). Modulatory role of dietary polyunsaturated fatty acids in Nrf2-mediated redox homeostasis. Progress in Lipid Research, 80, 101066.

29. Basil, M. C., & Levy, B. D. (2016). Specialized pro-resolving mediators: endogenous regulators of infection and inflammation. Nature Reviews Immunology, 16(1), 51–67.

30. Azizpor, P., Okakpu, O. K., Parks, S. C., Chavez, D., Eyabi, F., Martinez-Beltran, S., Nguyen, S., & Dillman, A. R. (2024). Polyunsaturated fatty acids stimulate immunity and eicosanoid production in Drosophila melanogaster. Journal of Lipid Research, 65(9), 100608.

31. Panettieri, S., Paddibhatla, I., Chou, J., Rajwani, R., Moore, R. S., Goncharuk, T., John, G., & Govind, S. (2020). Discovery of aspirin-triggered eicosanoid-like mediators in a *Drosophila* metainflammation blood tumor model. Journal of Cell Science, 133(5), jcs236141.

32. Yu, X., Song, S., Li, Z., Wang, T., Huang, H., Shen, Q., Wu, Z., & Luo, P. (2025). Targeted LC-MS/MS method of oxylipin profiling reveals differentially expressed serum metabolites in type 2 diabetes mice with panaxynol. Journal of Pharmaceutical and Biomedical Analysis, 253, 116540.

33. Reed, L. K., Williams, S., Springston, M., Brown, J., Freeman, K., DesRoches, C. E., Sokolowski, M. B., & Gibson, G. (2010). Genotype-by-Diet Interactions Drive Metabolic Phenotype Variation in *Drosophila melanogaster*. Genetics, 185(3), 1009–1019.

34. Dew-Budd, K., Jarnigan, J., & Reed, L. K. (2016). Genetic and Sex-Specific Transgenerational Effects of a High Fat Diet in Drosophila melanogaster. PLoS ONE, 11(8), e0160857.

35. Philippakis, A. A., Busser, B. W., Gisselbrecht, S. S., He, F. S., Estrada, B., Michelson, A. M., & Bulyk, M. L. (2006). Expression-Guided In Silico Evaluation of Candidate Cis Regulatory Codes for Drosophila Muscle Founder Cells. PLoS Computational Biology, 2(5), e53.

36. Maitra, U., Harding, T., Liang, Q., & Ciesla, L. (2021). GardeninA confers neuroprotection against environmental toxin in a Drosophila model of Parkinson’s disease. Communications Biology, 4(1), 162.

37. Chi, W., Iyengar, A. S., Fu, W., Liu, W., Berg, A. E., Wu, C. F., & Zhuang, X. (2022). Drosophila carrying epilepsy-associated variants in the vitamin B6 metabolism gene PNPO display allele-and diet-dependent phenotypes. Proceedings of the National Academy of Sciences, 119(9), e2115524119.

38. Kasuya, J., Johnson, W., Chen, H. L., & Kitamoto, T. (2023). Dietary supplementation with milk lipids leads to suppression of developmental and behavioral phenotypes of hyperexcitable Drosophila mutants. Neuroscience, 520, 1–17.

39. Kumar, R., Fatima, Z., Kumar, P., Kumar, P., Chauhan, B. S., & Srikrishna, S. (2024). Untargeted massspectrometry based lipidomics analysis reveals altered lipid profiles in a scribble knockdown-induced colorectal cancer model of Drosophila.

40. Cetraro, N., & Yew, J. Y. (2022). In situ lipid profiling of insect pheromone glands by direct analysis in real time mass spectrometry. Analyst, 147(14), 3276–3284.

41. Maitra, U., Conger, J., Owens, M. M. M., & Ciesla, L. (2023). Predicting structural features of selected flavonoids responsible for neuroprotection in a Drosophila model of Parkinson’s disease. Neurotoxicology, 96, 1–12.

42. Youdim, K. A., Shukitt-Hale, B., & Joseph, J. A. (2004). Flavonoids and the brain: interactions at the blood–brain barrier and their physiological effects on the central nervous system. Free Radical Biology and Medicine, 37(11), 1683–1693.

43. McDonald, H., Li, Q., Ashaduzzaman, M., Zhao, C., Pan, S., Szulczewski, G. J., & Liang, Q. (2024). Quantitative MALDI-MS and Imaging of Fungicide Pyrimethanil in Strawberries with 2-Nitrophloroglucinol as an Effective Matrix. Journal of the American Society for Mass Spectrometry, 35(6), 1272–1281.

44. Kumar, D., Sinha, S. N., & Gouda, B. (2024). Novel LC-MS/MS method for simultaneous determination of monoamine neurotransmitters and metabolites in human samples. Journal of the American Society for Mass Spectrometry, 35(4), 663–673.

45. Drouin-Ouellet, J., St-Amour, I., Saint-Pierre, M., Lamontagne-Proulx, J., Kriz, J., Barker, R. A., & Cicchetti, F. (2015). Toll-like receptor expression in the blood and brain of patients and a mouse model of Parkinson’s disease. International Journal of Neuropsychopharmacology, 18(6).

46. Takano, T., Fiore, S., Maddox, J. F., Brady, H. R., Petasis, N. A., & Serhan, C. N. (1997). Aspirin-triggered 15-epi-lipoxin A4 (LXA4) and LXA4 stable analogues are potent inhibitors of acute inflammation: evidence for anti-inflammatory receptors. The Journal of experimental medicine, 185(9), 1693–1704.

47. Serhan, C. N., & Chiang, N. (2002). Lipid-derived mediators in endogenous anti-inflammation and resolution: lipoxins and aspirin-triggered 15-epi-lipoxins. The Scientific World Journal, 2(1), 169–204.

48. Dalli, J., Ramon, S., Norris, P. C., Colas, R. A., & Serhan, C. N. (2015). Novel proresolving and tissue-regenerative resolvin and protectin sulfido-conjugated pathways. The FASEB Journal, 29(5), 2120.

49. Recchiuti, A., & Serhan, C. N. (2012). Pro-resolving lipid mediators (SPMs) and their actions in regulating miRNA in novel resolution circuits in inflammation. Frontiers in immunology, 3, 298.

50. Serhan, C. N., Bäck, M., Chiurchiù, V., Hersberger, M., Mittendorfer, B., Calder, P. C., Waitzberg, D. L., Stoppe, C., Klek, S., Martindale, R. G., & the International Lipids in Parenteral Nutrition Summit 2022 Experts. (2024). Expert consensus report on lipid mediators: Role in resolution of inflammation and muscle preservation. The FASEB Journal, 38(10), e23699.

51. Fantin, M., Garelli, F., Napoli, B., Forgiarini, A., Gumeni, S., De Martin, S., Montopoli, M., Vantaggiato, C., & Orso, G. (2019). Flavonoids Regulate Lipid Droplets Biogenesis in *Drosophila melanogaster*. Natural Product Communications, 14(5), 1934578X19852430.

52. Hammad, L. A., Cooper, B. S., Fisher, N. P., Montooth, K. L., & Karty, J. A. (2011). Profiling and quantification of Drosophila melanogaster lipids using liquid chromatography/mass spectrometry. Rapid communications in mass spectrometry, 25(19), 2959–2968.

53. Balvers, M. G., Verhoeckx, K. C., Bijlsma, S., Rubingh, C. M., Meijerink, J., Wortelboer, H. M., & Witkamp, R. F. (2012). Fish oil and inflammatory status alter the n-3 to n-6 balance of the endocannabinoid and oxylipin metabolomes in mouse plasma and tissues. Metabolomics, 8(6), 1130–1147.

54. Miller, J. S., Nguyen, T., & Stanley-Samuelson, D. W. (1994). Eicosanoids mediate insect nodulation responses to bacterial infections. Proceedings of the National Academy of Sciences, 91(26), 12418–12422.

55. Dalli, J., Chiang, N., & Serhan, C. N. (2014). Identification of 14-series sulfido-conjugated mediators that promote resolution of infection and organ protection. Proceedings of the National Academy of Sciences, 111(44), E4753–E4761.

56. Dalli, J., Zhu, M., Vlasenko, N. A., Deng, B., Haeggström, J. Z., Petasis, N. A., & Serhan, C. N. (2013). The novel 13S, 14S-epoxy-maresin is converted by human macrophages to maresin 1 (MaR1), inhibits leukotriene A4 hydrolase (LTA4H), and shifts macrophage phenotype. The FASEB Journal, 27(7), 2573.

57. Dalli, J., & Serhan, C. N. (2012). Specific lipid mediator signatures of human phagocytes: microparticles stimulate macrophage efferocytosis and pro-resolving mediators. Blood, The Journal of the American Society of Hematology, 120(15), e60–e72.

58. Martin, Marie, Emie Debenay, Jeanne Bardinet, Adrien Peltier, Line Pourtau, David Gaudout, Sophie Layé, Véronique Pallet, Anne-Laure Dinel, and Corinne Joffre. “Plant extracts and omega-3 supplementation modulate hippocampal oxylipin profile in response to LPS-induced neuroinflammation.” Inflammation Research 73, no. 11 (2024): 2023–2042.

59. Spencer, J. P. (2010). Beyond antioxidants: the cellular and molecular interactions of flavonoids and how these underpin their actions on the brain. Proceedings of the Nutrition Society, 69(2), 244–260.

60. Kennedy, D. O., & Wightman, E. L. (2011). Herbal extracts and phytochemicals: plant secondary metabolites and the enhancement of human brain function. Advances in Nutrition, 2(1), 32–50.

61. Schaefer, S., Baum, M., Eisenbrand, G., Dietrich, H., Will, F., & Janzowski, C. (2006). Polyphenolic apple juice extracts and their major constituents reduce oxidative damage in human colon cell lines. Molecular nutrition & food research, 50(1), 24–33.

62. Toppo, E., Darvin, S. S., Esakkimuthu, S., Stalin, A., Balakrishna, K., Sivasankaran, K., Pandikumar, P., Ignacimuthu, S., & Al-Dhabi, N. A. (2017). Antihyperlipidemic and hepatoprotective effects of Gardenin A in cellular and high fat diet fed rodent models. Chemico-Biological Interactions, 269, 9–17.

63. Strathearn, K. E., Yousef, G. G., Grace, M. H., Roy, S. L., Tambe, M. A., Ferruzzi, M. G., Wu, Q.-L., Simon, J. E., Lila, M. A., & Rochet, J.-C. (2014). Neuroprotective effects of anthocyanin- and proanthocyanidin-rich extracts in cellular models of Parkinson’s disease. Brain Research, 1555, 60–77.

64. Seigler, D. S., Friesen, J. B., Bisson, J., Graham, J. G., Bedran-Russo, A., McAlpine, J. B., & Pauli, G. F. (2021). Do certain flavonoid IMPS have a vital function? Frontiers in Nutrition, 8, 762753.

65. Maitra, U., Stephen, C., & Ciesla, L. M. (2022). Drug discovery from natural products–Old problems and novel solutions for the treatment of neurodegenerative diseases. Journal of Pharmaceutical and Biomedical Analysis, 210, 114553.

66. Miller, S. J., Darji, R. Y., Walaieh, S., Lewis, J. A., & Logan, R. (2023). Senolytic and senomorphic secondary metabolites as therapeutic agents in Drosophila melanogaster models of Parkinson’s disease. Frontiers in Neurology, 14, 1271941.

67. Hack, W., Gladen-Kolarsky, N., Chatterjee, S., Liang, Q., Maitra, U., Ciesla, L., & Gray, N. E. (2024). Gardenin A treatment attenuates inflammatory markers, synuclein pathology and deficits in tyrosine hydroxylase expression and improves cognitive and motor function in A53T-α-syn mice. Biomedicine & Pharmacotherapy, 173, 116370.

68. Ferger, B., Teismann, P., Earl, C. D., Kuschinsky, K., & Oertel, W. H. (1999). Salicylate protects against MPTP-induced impairments in dopaminergic neurotransmission at the striatal and nigral level in mice. Naunyn-Schmiedeberg’s archives of pharmacology, 360, 256–261.

69. Abohassan, M., Fayzullaev, N., Edris, G., Uthirapathy, S., Sanghvi, G., Vakada, N. B. R., Sharma, S., Nakash, P., Mustafa, Y. F., & Shallan, M. A. (2025). Supercritical extraction of salicin, aspirin precursor, from the willow bark, laboratory optimization via response surface methodology and mathematical modeling. The Journal of Supercritical Fluids, 219, 106540.

70. Navale, S. S., Mulugeta, A., Zhou, A., Llewellyn, D. J., & Hyppönen, E. (2022). Vitamin D and brain health: an observational and Mendelian randomization study. The American journal of clinical nutrition, 116(2), 531–540.

71. Michelakos, T., Kousoulis, A. A., Katsiardanis, K., Dessypris, N., Anastasiou, A., Katsiardani, K.-P., Kanavidis, P., Stefanadis, C., Papadopoulos, F. C., & Petridou, E. Th. (2013). Serum Folate and B12 Levels in Association With Cognitive Impairment Among Seniors: Results From the VELESTINO Study in Greece and Meta-Analysis. Journal of Aging and Health, 25(4), 589–616.

72. Yang, S. et al. Curcumin protects dopaminergic neuron against LPS induced neurotoxicity in primary rat neuron/glia culture. Neurochem. Res. 33, 2044–2053 (2008).

73. Kretzer, C., Jordan, P. M., Meyer, K. P. L., Hoff, D., Werner, M., Hofstetter, R. K., Koeberle, A., Cala Peralta, A., Viault, G., Seraphin, D., Richomme, P., Helesbeux, J.-J., Stuppner, H., Temml, V., Schuster, D., & Werz, O. (2022). Natural chalcones elicit formation of specialized pro-resolving mediators and related 15-lipoxygenase products in human macrophages. Biochemical Pharmacology, 195, 114825.

74. Tamtaji, O. R., Taghizadeh, M., Aghadavod, E., Mafi, A., Dadgostar, E., Daneshvar Kakhaki, R., Abolhassani, J., & Asemi, Z. (2019). The effects of omega-3 fatty acids and vitamin E co-supplementation on gene expression related to inflammation, insulin and lipid in patients with Parkinson’s disease: A randomized, double-blind, placebo-controlled trial. Clinical Neurology and Neurosurgery, 176, 116–121.

75. Werz, O., Gerstmeier, J., Libreros, S. et al. Human macrophages differentially produce specific resolvin or leukotriene signals that depend on bacterial pathogenicity. Nat Commun 9, 59 (2018).

76. Bischoff-Ferrari, H.A., Gängler, S., Wieczorek, M. et al. Individual and additive effects of vitamin D, omega-3 and exercise on DNA methylation clocks of biological aging in older adults from the DO-HEALTH trial. Nat Aging 5, 376–385 (2025).

77. Wooten, G. F., Currie, L. J., Bovbjerg, V. E., Lee, J. K., & Patrie, J. (2004). Are men at greater risk for Parkinson’s disease than women? Journal of Neurology, Neurosurgery & Psychiatry, 75(4), 637–639.

78. J. Lee, P. Pinares-Garcia, H. Loke, S. Ham, E. Vilain, & V.R. Harley (2019). Sex-specific neuroprotection by inhibition of the Y-chromosome gene, *SRY*, in experimental Parkinson’s disease, Proc. Natl. Acad. Sci. U.S.A. 116 (33) 16577–16582.

79. Westfall, S., Lomis, N., & Prakash, S. (2019). A novel synbiotic delays Alzheimer’s disease onset via combinatorial gut-brain-axis signaling in Drosophila melanogaster. PLoS One, 14(4).

80. Jourová, L., Vavreckova, M., Zemanova, N., Anzenbacher, P., Langova, K., Hermanova, P., Hudcovic, T., & Anzenbacherova, E. (2020). Gut Microbiome Alters the Activity of Liver Cytochromes P450 in Mice With Sex-Dependent Differences. Frontiers in Pharmacology, 11, 01303.

