## Supplementary material for "Plant-derived soft electrophiles upregulate pro-resolving oxylipins in a paraquat-induced *Drosophila* model of Parkinson’s disease": APPENDIX A - Supplementary information.docx

**APPENDIX A - SUPPLEMENTAL INFORMATION**

1. **Chromatogram representing the detection of 13-HODE and 13-oxoODE in the fly heads (Linoleic acid+GardeninA > PQ in comparison to Standard fly food > PQ).**

**[A] (Linoleic acid + GardeninA) > PQ in *Canton S* adult male heads**

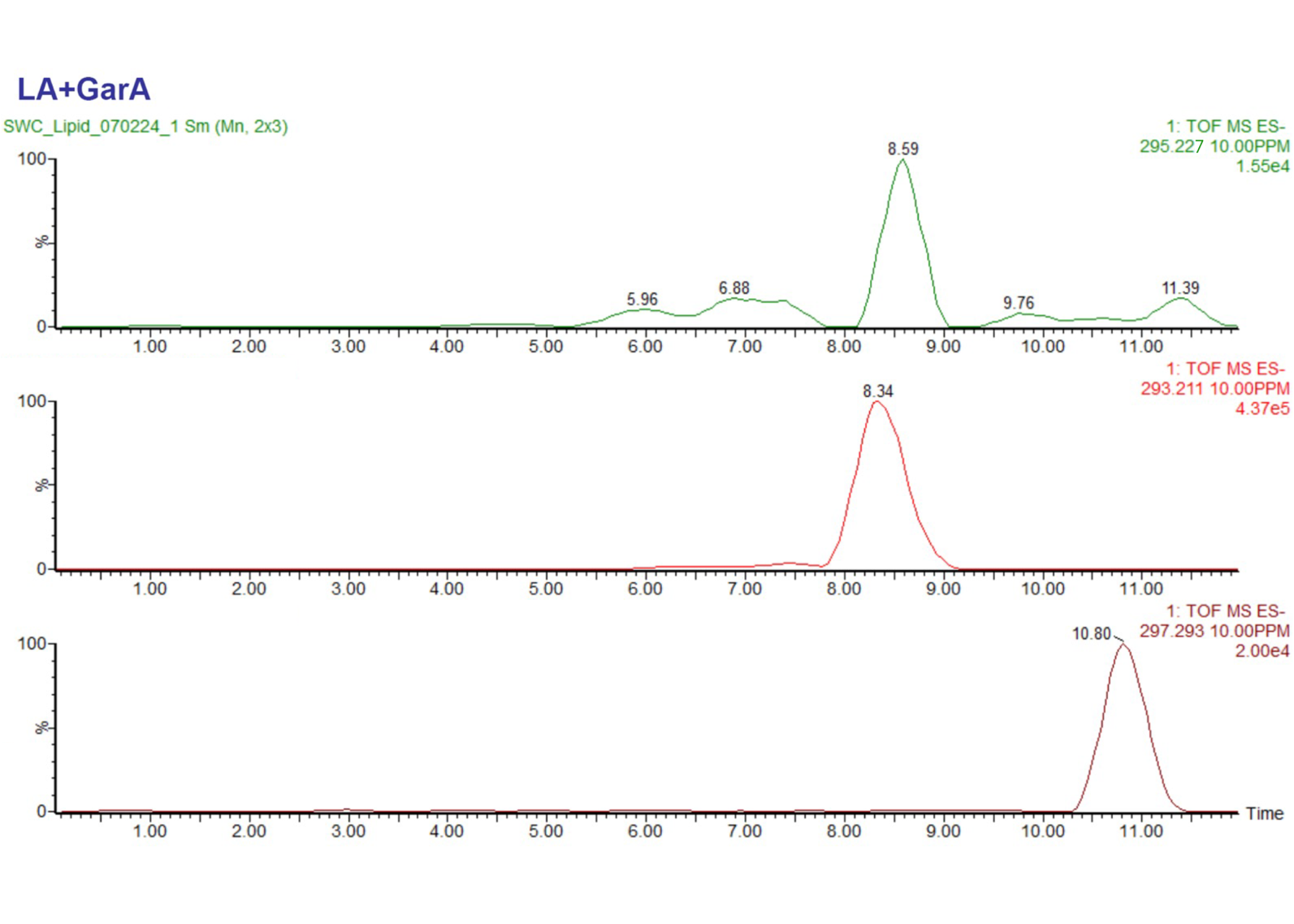

^13^C_18_ Linoleic acid

Internal standard

297.2928 m/z

13-oxoODE

293.2117 m/z

13-HODE

295.2273 m/z

**[B] Standard fly food > PQ in *Canton S* adult male heads**

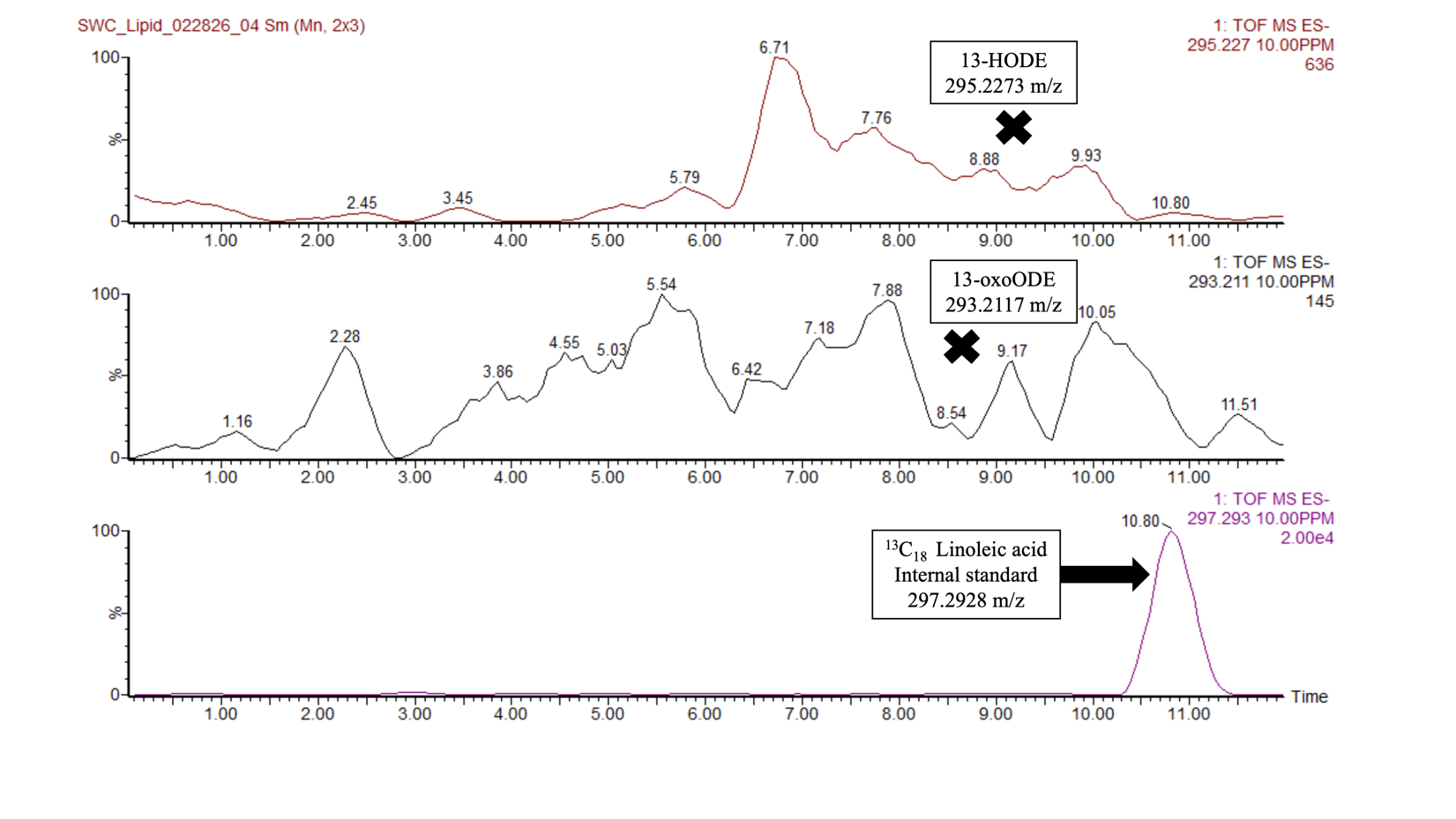

2. **Verification of LOD and LOQ calculation:**

LOD (Limit of detection) was found to be 0.39 nmol/mL (0.12 ppm) for 13-HODE and 0.36 nmol/mL (0.11ppm) for 13-oxoODE, and the limit of quantification LOQ (Limit of Quantitation) was found to be at 1.21 nmol/mL (0.35 ppm) for 13-HODE and 1.13 nmol/mL (0.33 ppm) for 13-oxoODE**.** For calculation purposes, the untreated (flies only kept in standard diet) fly head homogenate matrix is treated as the analyte blank. The following calibration curves were used for the calculation of LOD and LOQ**:**

**[A]** **For 13-oxoODE:**

Calibration curve for LOD/LOQ Calculation: 13-oxoODE

| Concentration (in nmol/mL | 13-oxoODE/IS Peak area ratio |
| --- | --- |
| 1.05 | 1.07511 |
| 4.22 | 1.0711 |
| 8.43 | 1.0815 |
| 16.87 | 1.10181 |
| 33.73 | 1.13753 |

Calculated standard deviation of blank (σ) = 0.00023

Slope of the calibration curve (S) = 0.0021

LOD = [3.3 * σ] / S = [3.3 * 0.00023] / 0.0021 = 0.36 nmol/mL (~0.11 ppm)

LOQ = [10 * σ] / S = [10 * 0.00023] / 0.0021 = 1.13 nmol/mL (~0.33 ppm)

**[B] For 13-HODE:**

Calibration curve for LOD/LOQ Calculation: 13-HODE

| Concentration (in nmol/mL) | 13-HODE/IS Peak area ratio |
| --- | --- |
| 1.05 | 1.05511 |
| 4.22 | 1.0671 |
| 8.43 | 1.0725 |
| 16.87 | 1.09181 |
| 33.73 | 1.11753 |

Calculated standard deviation of blank (σ) = 0.00023

Slope of the calibration curve (S) = 0.0019

LOD = [3.3 * σ] / S = [3.3 * 0.00023] / 0.0019 = 0.39 nmol/mL (~0.12 ppm)

LOQ = [10 * σ] / S = [10 * 0.00023] / 0.0019 = 1.21 nmol/mL (~0.36 ppm)

3. **Verification of food uptake**

The amount of food consumed by the flies is the basis for all feeding assays and the detection of oxylipins post-PQ exposure. Especially in this case, the supply of Linoleic acid was mandatory to ensure the production of oxylipins, because linoleic acid is not produced by the flies themselves and is also not present in the composition of the standard fly food. Being an essential omega-6-derived fatty acid, it needs to be acquired only through diet. However, the taste and odor of various phytochemicals and linoleic acid combined with the standard fly food may affect feeding behavior and complicate the interpretation of treatment effects. Additionally, linoleic acid was utilized to enrich the standard fly meal in both the sample and control groups. Hence, food absorption was confirmed by employing UPLC-HRMS to identify a linoleic acid peak in fly head homogenates. The chromatogram below shows the detection of the linoleic acid peak in the control fly head homogenate, where the flies were kept in the standard fly food supplemented with Linoleic acid for 5 days and then exposed to PQ. The presence of the linoleic acid peak confirms that the food uptake was optimal. Peaks for 13-oxoODE and 13-HODE are absent in the chromatogram below, which indicates that the detection and quantification of these oxylipins as represented in our other chromatograms, is possible due to upregulation by our dietary soft electrophiles and not just linoleic acid. When taken as a whole, these results show similar food consumption levels in all treatment groups, suggesting that the observed experimental effects are unlikely to be due to changed feeding behavior, metabolism, or sexual dimorphism.

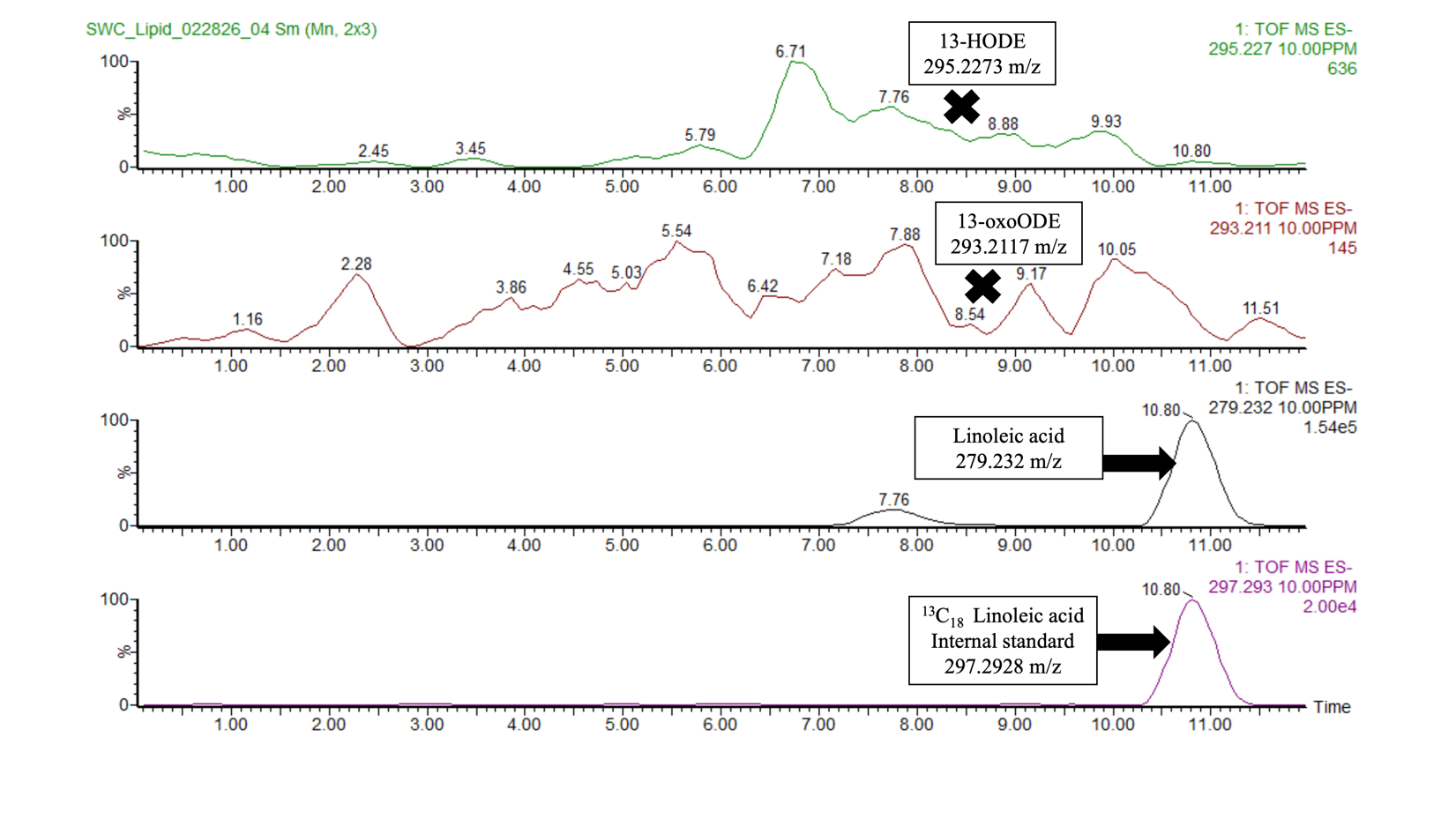

4. **Structures representing the group of flavonoids that did not show any effect on the lifespan and climbing index post-PQ exposure in the wild-type Canton S male/female flies.** Here the structures of specific flavonoids that did not show any protective activity is compared to that of Gardenin A which has previously been reported as neuroprotective in both *Drosophila* [36] as well as mice [67].

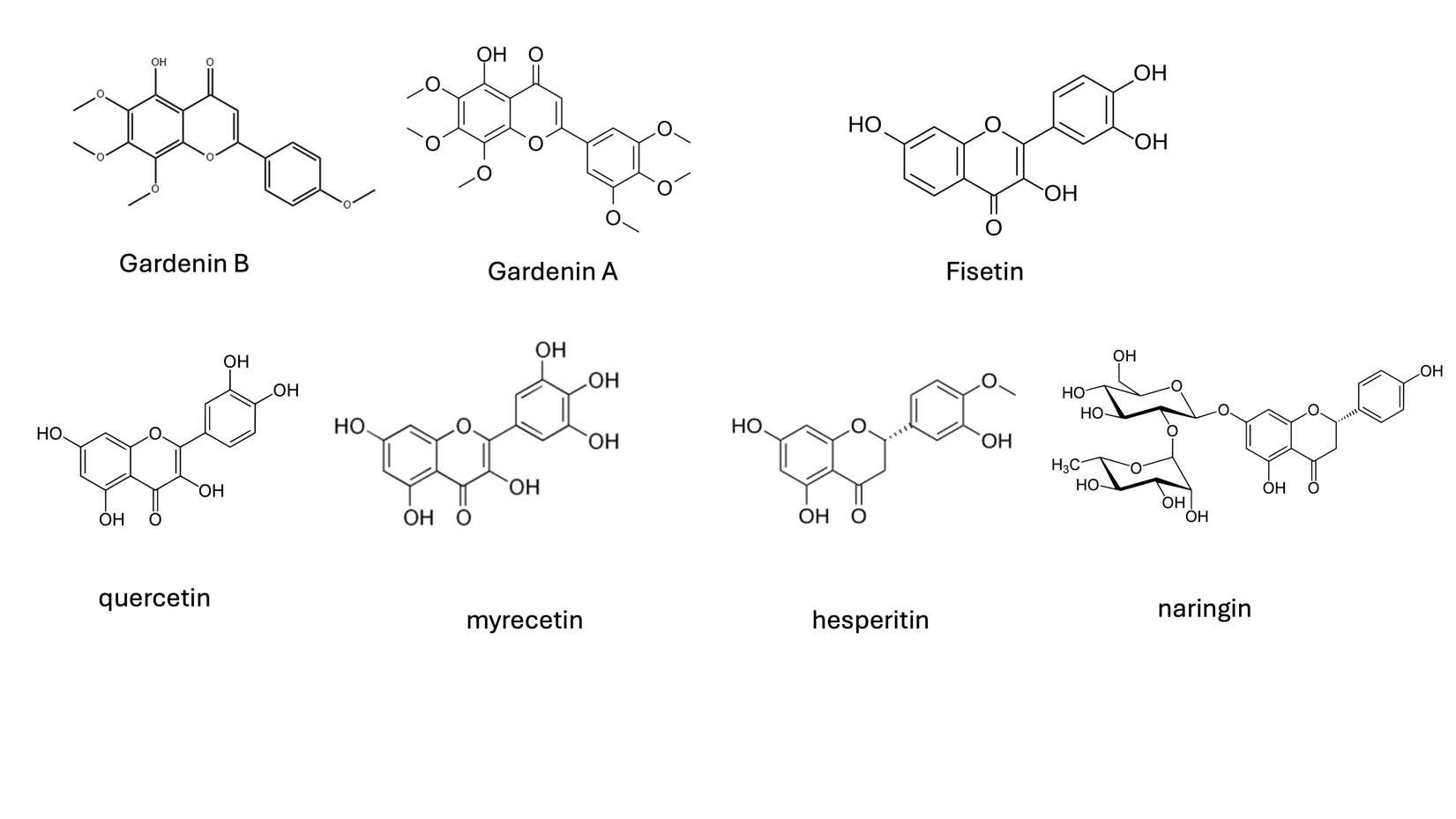
